# Systematic functional screening of chromatin factors identifies strong lineage and disease dependencies in normal and malignant haematopoiesis

**DOI:** 10.1101/2022.08.11.503571

**Authors:** D Lara-Astiaso, A Goñi-Salaverri, J Mendieta-Esteban, N Narayan, C Del Valle, T Gross, G Giotopoulos, M Navarro-Alonso, J Zazpe, F Marchese, N Torrea, IA Calvo, C Lopez, D Alignani, A Lopez, B Saez, J. P Taylor-King, F Prosper, N Fortelny, B. J. P Huntly

## Abstract

Interactions between transcription factors (TF) and chromatin factors (CF) regulate gene expression programmes to determine cellular fate. However, unlike for TF, the exact role of CF in this process is poorly understood. Using haematopoiesis as a model system and utilising novel functional CRISPR screens *ex vivo* and *in vivo*, coupled with Perturb-Seq, CF binding and genome-wide chromatin accessibility in primary murine cells, we assess the role of 550 chromatin factors in lineage choice in normal haematopoiesis and the maintenance of acute myeloid leukaemia (AML). These studies demonstrate marked specificity for a large number of CFs in lineage determination, highlighting functional diversity within specific families of chromatin regulators, including MLL-H3K4-methyltransferases and different BAF-complexes, that regulate disparate lineage decisions across haematopoiesis. Conversely, we demonstrate that unrelated Repressive complexes function similarly to restrain excessive myeloid differentiation and protect lineage diversity. We identify interactions between CF and TF that, at least in part, explain the regulatory function of CF and link *Brd9-*loss to a premalignant state. Utilising similar experiments in a relevant murine AML model, we demonstrate opposing effects for CF in normal haematopoiesis and AML, where MLL-H3K4-methyltransferases, c-BAF-remodelers and Repressive complexes prevent differentiation and maintain leukaemic fitness. We show that this alteration relates to differential utilisation of TF by CF complexes between normal and malignant haematopoiesis, highlighting corrupted TF-CF interactions as potential novel avenues for therapeutic intervention in AML. Together, this study provides novel insights on the functional diversity of chromatin factors in governing cell-fate.

## Introduction

Cell-fate decisions are governed by the coordinated activities of Transcription (TFs) and Chromatin Factors (CFs) that together form gene regulatory complexes (GRC) to orchestrate tissue-specific transcriptional patterns and drive cellular phenotype^1^. The widescale description of TF binding and its relationship to chromatin accessibility and gene expression, obtained from ChIP-Seq, ATAC-Seq and transcriptional “maps” derived across developmental processes, have provided us with a highly developed understanding of the instructional role for TF in governing cell-fate^2,3^. Conversely, although the role of individual CF, particularly those mutated across malignancies^4^, are being elucidated, we still lack a proper understanding of CF functions in differentiation. Specifically, whether CF have specific or redundant roles during lineage differentiation, how they functionally interact with individual TFs and the identity of these TF-CF interactions remain unresolved questions.

We have chosen to address these fundamental questions in the exemplar process of haematopoiesis, where multiple mature cells with specific functions (oxygen carriage, formation of blood clots, fighting infections, etc.) derive from a single self-renewing cell type, the hematopoietic stem cell (HSC)^5^. The study of haematopoiesis benefits from a comprehensive cellular blueprint^3^ that describes normal differentiation and malignant transformation and a well-annotated molecular blueprint^2,3,4^, including a detailed transcriptional landscape with single cell resolution and comprehensive mapping of TF activity across the haematopoietic lineages derived from ATAC-seq, ChIP-seq and knockout mouse models^7,8-12^. In addition, the importance of both CF and TF has been further emphasised by studies conducted over the last decade that have documented mutations altering the function of TF and CF to be highly recurrent and almost uniform in haematological malignancies such as acute myeloid leukaemia (AML)^13^.

In this report, we have functionally characterised the requirement of 550 CFs in normal lineage differentiation and further dissected, with single-cell resolution, the roles of 40 CFs during *in vivo* lineage commitment (Perturb-seq). Using epigenetic profiling (ATAC-seq and ChIP-seq) we probe TF-CF interactions with lineage regulatory potential. Finally, we compare the functions of key CFs and their TF-associations between normal and malignant haematopoiesis. These studies demonstrate, both *ex vivo* and *in vivo*, that similar complexes, including three MLL-H3K4 COMPASS methyltransferases (KMTs) and non-canonical (nc-BAF) and canonical BAF (c-BAF) complexes, regulate disparate lineage decisions across haematopoiesis. However, somewhat paradoxically, we demonstrate that unrelated Repressive complexes function similarly, but non-redundantly, to restrain excessive myeloid differentiation and protect lineage diversity. We identify interactions between CF and TF that explain the regulatory function of CF and demonstrate that KO of the nc-BAF member *Brd9* leads to a premalignant accumulation of myeloid progenitors related to impaired recruitment of late myeloid TFs. Finally, utilising similar experiments in a relevant murine AML model, we describe differential utilisation of TF by CF complexes between normal and malignant haematopoiesis. Moreover, we identify perturbations of MLL-, BAF- and Repressive CF that trigger partial differentiation and loss of leukaemic potential and demonstrate maintenance of this potential through corrupted Stat5a-CF interactions, thereby identifying known and potential novel targets for therapeutic intervention in AML. Taken all together, our work greatly elucidates the role of CF and their TF partners across normal and malignant haematopoiesis.

## Results

### Functional screens identify linage specificities for Chromatin Factors in haematopoiesis

To interrogate the roles of CFs in regulating normal haematopoiesis, we developed 4 screening platforms that couple cytokine-instructed differentiation of primary haematopoietic progenitors with FACS readouts to study 4 key lineage transitions (Fig 1a-b): **1)** The balance between self-renewal and differentiation, **2)** early branching into myeloid and mega-erythroid lineages **3)** full differentiation of multipotent progenitors into myeloid fate (spanning both early and late transitions) and **4)** full differentiation of myeloid primed progenitors (GMPs) into mature myeloid cells (only late transitions). To validate our screens, we cross-referenced the expression profiles of each readout population to bulk and single-cell expression maps of haematopoiesis and demonstrated fidelity with their *in vivo* counterparts (Fig S1a-b). Next, we generated a CRISPR library targeting 550 genes that included every CF expressed by myeloid and mega-erythroid lineages *in vivo* (Table S1). Then, we delivered our library to both Cas9 (GFP+) and Non-Cas9 (GFP-) progenitors, co-cultured them throughout our 4 differentiation conditions and used surface differentiation markers to evaluate the sgRNA distributions in the different readout populations emerging from Cas9 or Non-Cas9 progenitors, separated by GFP surface expression. The inclusion of Non-Cas9 cells allowed us to account for potential biases in the CRISPR library distribution related to clonal expansion. This analysis showed differential enrichment between readout populations for the Cas9 progenitors, but no difference between Non-Cas9 progenitors ruling out potential biases in the CRISPR library distribution (Fig S1c). Finally, using the Non-Cas9 normal distributions as a background reference, we calculated a lineage score for each CF that represented their contribution to each of the 4 lineage transitions (Fig S1d)

**Figure 1.**
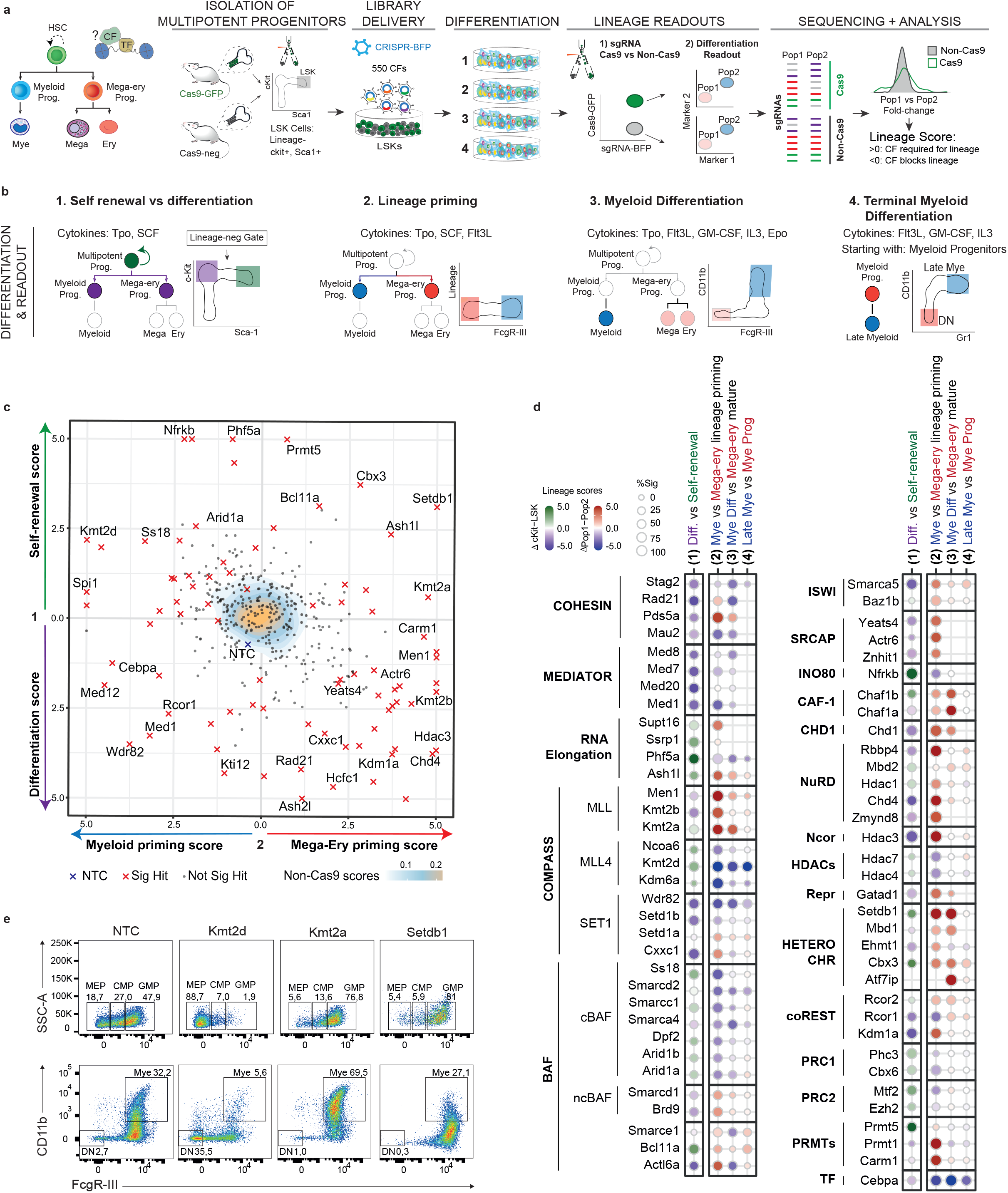
Functional interrogation of Chromatin Factors (CF) reveals strong lineage dependencies during haematopoietic differentiation. (**a**) Schematic drawing of the CRISPR screen workflow **(left to right)-** <u>Isolation of Multipotent progenitors</u> (Lin-, Sca1+, ckit+) from Cas9-GFP and Non-Cas9 mouse strains; <u>Delivery of the CRISPR library</u> targeting 550 Chromatin Factors to the mixed Cas9 and non-Cas9 population of multipotent progenitors; <u>Differentiation</u> of CRISPR modified multipotent progenitors using 4 *ex vivo* culture systems (details in panel 1b); <u>FACS-based differentiation readouts</u> (see panel 1b) from Cas9 and Non-Cas9 fractions; <u>Quantification of sgRNA distributions</u> across the readout populations; <u>Calculation of lineage scores</u> for each Chromatin Factor, using the Non-Cas9 sgRNA distributions as a background model to correct for library bias and dispersion. **(b)** Differentiation systems and FACS-based Readouts. <u>1- Self-Renewal vs Differentiation</u>: culture driven by SCF and Tpo where Multipotent (Lin-, ckit+, Sca1+) and Differentiated (Lin-, ckit+, Sca1-) populations are evaluated. <u>2- Lineage priming</u>: culture driven by SCF, Flt3L, and Tpo where Mega-erythroid (Lin-, ckit+, Sca1-, FcgR- III-) and Myeloid (Lin-, ckit+, Sca1-, FcgR-III+) progenitors are evaluated. 3<u>- Myeloid Differentiation</u>: culture driven by GM-CSF, G-CSF, Flt3L, IL3 and EPO where Mature myeloid (FcgR-III+, CD11b+) and Non-Myeloid (FcgR-III-, CD11b-) lineages are evaluated. 4<u>- Terminal Myeloid Maturation</u>: culture driven by GM-CSF, G-CSF, Flt3L, IL3 where Immature myeloid (CD11b-, Gr1-) and Mature myeloid (CD11b+, Gr1+) lineages are evaluated. Screens 1-3 are initiated from LSK and screen 4 from committed myeloid progenitors (Granulocyte macrophage progenitors, GMPs). **(c)** 2D Projection of lineage scores for “1- Differentiation vs Self-renewal” (y-axis) and “2- Lineage priming” into myeloid vs mega-erythroid (x-axis). For each gene, lineage scores are shown aggregated across all guides and libraries. Genes with significant changes are shown with a red cross. Genes that did not reach significance thresholds are shown as black dots, aggregated NTCs are shown with a blue cross. Data for Non-Cas9 cells are shown in the background using a yellow-blue density. (**d**) Lineage scores for chromatin factors grouped on the basis of complex membership. The colour of each dot represents the lineage score. The size of the dot represents the number of significant guides. (**e**) Exemplar immunophenotypic validations for CFs with strong scores in the lineage priming (top) and myeloid differentiation (bottom) systems.

We identified 121 CFs with significant lineage scores that span diverse roles in the differentiation processes and often demonstrated antagonist behaviours (Fig 1c, S1e). Since CF work in the context of specific complexes, we grouped CFs based on known complex membership, revealing a high degree of phenocopy amongst well-characterised complex members (Fig 1d). For instance, while members of the RNA elongation machinery (*Pds5a, Ash1l*) preserve progenitor multipotency, members of the Cohesin and Mediator complexes are required for differentiation into both myeloid and mega-erythroid fates, confirming that chromatin looping is a general requirement for lineage progression^14,15^ (Fig 1c-d). Furthermore, the screens detected strong lineage specificities within and between complexes exerting related epigenetic activities (Fig S1e-f). The three different COMPASS H3K4 methylases^16^ demonstrated very different requirement patterns, with MLL1 members presenting a strong mega-erythroid dependency but MLL4 members, in particular the catalytic subunit *Kmt2d*, behaving as strong myeloid regulators required for both priming and later myeloid maturation. Lastly, similarly to Cohesin and Mediator, SET1 components were required to initiate differentiation but displayed antagonistic behaviours at lineage branching points, where *Wdr82* acted as a pro-myeloid factor, but *Cxxc1* and *Setd1a* demonstrated mega-erythroid dependency; suggesting the existence of functionally diverse SET1 subcomplexes.

Similarly, ATP-dependent chromatin remodelers presented distinct lineage dependencies. Loss-of-function of NuRD and ISWI, complexes predominantly mediating nucleosome sliding and repressive roles^17^, facilitated myeloid priming at the expense of mega-erythroid fates. Conversely, components of the BAF complex, which catalyse nucleosome eviction^18^, appear predominantly pro-myeloid regulators, with the exception of ncBAF subunits *Brd9* and *Smarcd1*, which exhibited weak mega-erythroid dependency. Finally, similarly to NuRD and ISWI, members of the SRCAP complex, which deposits H2A.Z^19^, showed a pro-mega/erythroid role, while *Nfrkb*, a component of INO80, which removes H2A.Z^20^, ranked as a top regulator of progenitor multipotency. Analysis of Repressive complexes demonstrated a degree of functional heterogeneity, however the strongest effects were observed for factors that collectively behaved as myeloid repressors including regulators of heterochromatin formation (*Setdb1, Cbx3*), histone deacetylases *Hdac1* and *Hdac3*, (member of the Ncor complex) and most components of the coREST corepressor complex, except *Rcor1*, especially its catalytic subunit the H3K4 demethylase *Kdm1a/Lsd1*^21^. The screen-based phenotypes of exemplar CF were confirmed in sgRNA analyses (Fig 1e, S2a-d).

Collectively, these findings highlight significant functional diversity within related activatory complexes (i.e COMPASS and BAF remodelers) suggesting that specific CF subcomplexes work at different stages along differentiation trajectories. By contrast, disparate complexes predominantly involved in epigenetic repression, like NuRD and ISWI remodelers or various corepressors, function in a more uniform manner to instruct mega-erythroid lineages, likely by repressing myeloid fates^22^.

### In vivo Perturb-seq highlights functional diversity for similar CF complexes during lineage specification

Next, we decided to explore the functional diversity of CFs in their proper physiological context, utilising Perturb-seq to profile the *in vivo* developmental patterns of 40 CF knockouts. These comprised COMPASS H3K4 methyltransferases, Chromatin Remodelers and Repressive Complexes that demonstrated strong lineage dependencies *ex vivo*, as well as factors with strong effects in regulating the self-renewal vs differentiation balance of early progenitors, such as the Arginine-methyl-transferase *Prmt5* or the Cohesin subunit *Stag2*.

Following CRISPR-KO of these CF in multipotent progenitors (LSKs), they were transplanted into sublethally irradiated mice and, at 2-weeks post-transplant, we profiled their progeny with Perturb-seq in Lineage- and Lineage+/c-kit+ fractions (Fig 2a). In combination, these two populations reconstituted the main lineage trajectories emerging from transplanted LSKs; myeloid, mega-erythroid, basophil and lymphoid fates, with most cells spanning myeloid and erythroid branches (Fig 2b-c, S3a-d). In addition, we complemented this approach with Perturb-seq profiling of *ex vivo* culture systems that examined lineage priming and myeloid-differentiation conditions (Fig S3e-f, S4b). Analysis of non-targeting control (NTC) guide distribution demonstrated a homogeneous distribution across all lineages, ruling out potential confounding sgRNA patterns arising from clonal selection (Fig 2d-e, S3c). Conversely, CF-KOs demonstrated striking lineage specific patterns, which were quantified by assessing the enrichment or depletion of each CF-KO across the main haematopoietic lineages and by measuring the transcriptional progression of CF-KOs along myeloid and erythroid differentiation trajectories using pseudotime analysis (Fig 2f-g, S4a, c-d).

**Figure 2.**
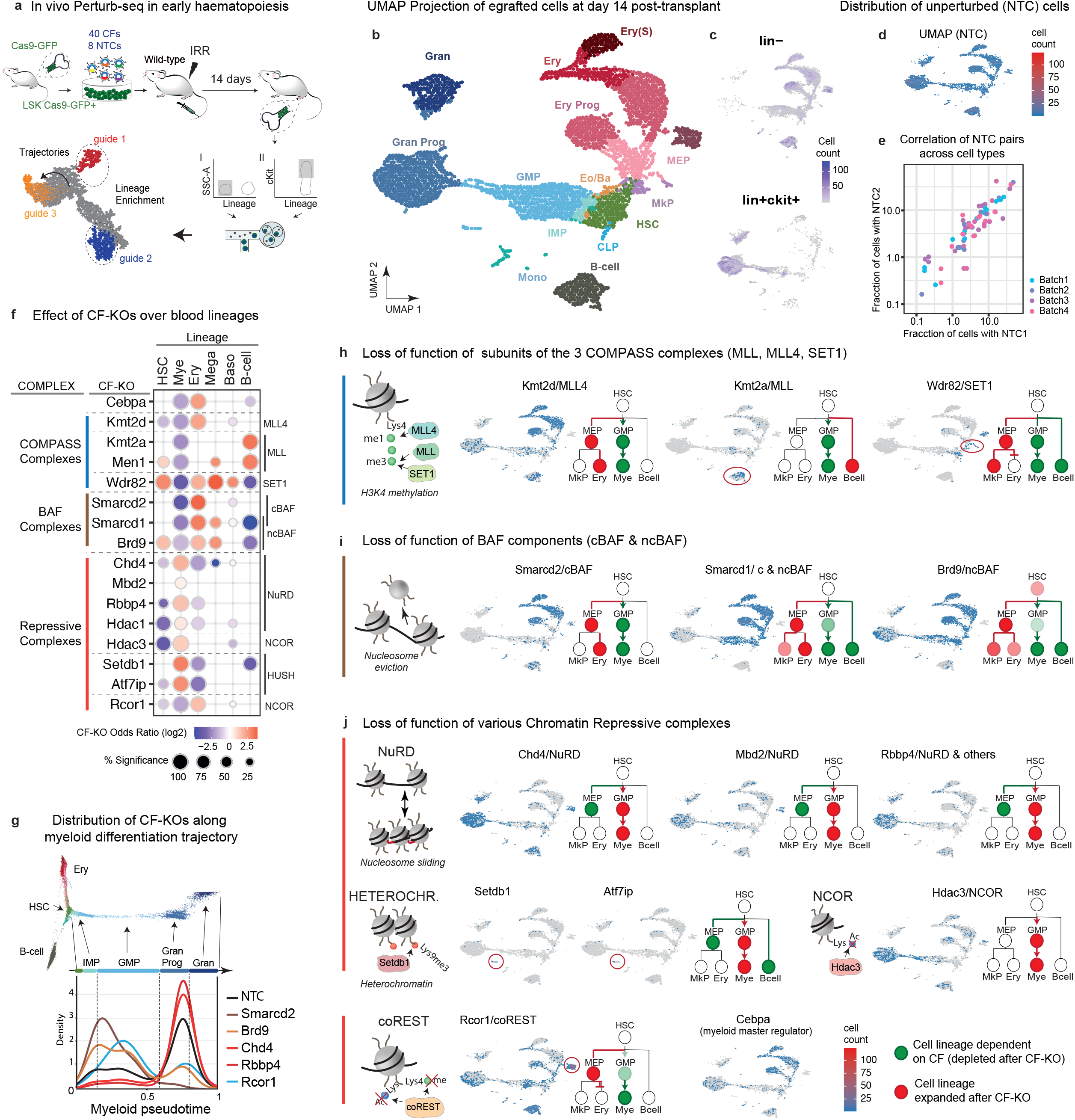
*In vivo* Perturb-seq highlights functional diversity for CF Complexes during lineage specification. (**a**) Schematic drawing of the *in vivo* Perturb-seq experiment. (**b-c**) Single-cell characterisation of the bone-marrow transplant. **(b)** UMAP projection of the single-cell transcriptomes derived from transplanted progenitors at day 14 post-transplant. The areas in the UMAP are coloured and named according to the predominant cell type. Cell clusters were annotated using external reference maps of haematopoiesis^48^. (**c**) UMAP projection of the Lineage- and Lineage+ ckit+ fractions. The colour scale represents the number of cells in each area. **(d-e)** Assessment of potential biases in the *in vivo* Perturb-seq system. **(d)** UMAP projection of the cells harbouring Non-targeting control (NTC) guides. **(e)** Scatterplot showing high correlation between pairs of individual NTC guides across all haematopoietic populations in 4 experimental batches. Each dot represents the abundance of NTC1 and NTC2 in a given population. The high correlation of NTC abundances validates the absence of clonal effects (**f-g**) Single-cell characterization of Chromatin Complex perturbation on haematopoietic differentiation. **(f)** Enrichment/Depletion analyses of CF-KOs across the main haematopoietic lineages: HSC (progenitor), Myeloid, Erythroid, Megakaryocyte, Basophil, B-cell. The Myeloid lineage comprises Immature Myeloid Progenitors, IMPs, GMPs, Monocytes, Granulocyte Progenitors and Granulocytes. The Erythroid lineage comprises MEPs, Erythroid Progenitors and Mature Erythroid cells. Dot colour and size relate to the log2 odds ratio and the percent of significant enrichments versus NTCs, respectively. **(g)** Trajectory analysis showing the evolution of specific CF-KOs along Myeloid differentiation. Cell are ordered from HSCs to mature granulocytes using pseudotime. The NTC distribution is represented with a black line. Individual KOs are colour coded as per the key on the right. The differentiation stages are colour-coded along the top of the graph from green (HSC) to dark blue (Mature Granulocytes) **(h-j)** Graphic representation of the roles key Chromatin Regulatory Complexes: (left) generic schematic of epigenetic activity for each complex; (mid) UMAP as per (**b**) showing CF-KO distributions, where cells are aggregated and the colour of each area represents the number of cells; (right) schematic haematopoietic tree representing the developmental effect of the CF-KO. (**h**) Phenotypes for key subunits belonging to the different COMPASS H3K4 methylation complexes. (**i**) Phenotypes for key subunits belonging to canonical and non-canonical BAF remodelers. Of note, Smarcd1 is shared between both complexes. (**j**) Phenotypes for key subunits belonging to NuRD Remodelers and other Repressive complexes.

Assessing KO of the three COMPASS H3K4 methyltransferase complexes generally confirmed the lineage dependencies found in the *ex vivo* screens for components of the MLL4 and SET1 COMPASS complexes (Fig 2f, 2h). Loss of function of the MLL4 catalytic subunit, *Kmt2d*, altered the myeloid vs erythroid balance by completely disrupting myeloid differentiation and accelerating erythroid lineage progression. Conversely, disruption of the SET1 complex, via *Wdr82*-KO, abolished both myeloid and lymphoid lineages and provoked stalled mega-erythroid trajectories, with *Wdr82*-KO cells accumulating at the multipotent (HSC) and especially megakaryocytic (Mkp) progenitor stages. In contrast to MLL4 and SET1, disruption of the MLL1 complex (*Kmt2a*-KO and *Men1*-KO) *in vivo* did not recapitulate the pattern of *ex vivo* loss (Fig 2f, 2h S4a, c); while the disruption of *Kmt2a* and *Men1* favoured granulocytic differentiation under strong *ex vivo* myelo-erythroid cytokine instruction, their KO *in vivo* promoted B-cell priming. Such distinct behaviour is likely due to the different signals encountered by the MLL1 complex-perturbed cells in both systems, highlighting that CF function can be regulated by extrinsic signals. These results demonstrate a lack of redundancy between the three writers of H3K4 methylation and suggest that cell-type specific H3K4me patterns are specifically deposited by particular epigenetic writers and are required at different hematopoietic stages.

Analysis of BAF perturbation *in vivo* confirmed myeloid dependency for the cBAF complex, with disruption of *Smarcd2* causing early erythroid skewing and accelerated erythropoiesis (Fig2f-g, i), a pattern previously reported in the *Smarcd2*-KO mouse strain^23^. In contrast, disruption of the ncBAF complex, defined by Brd9, abolished lymphoid development and mildly impeded myeloid priming (Fig 2f, 2i). Despite the modest myeloid defect observed for *Brd9*-KO cells, pseudotime analysis detected a marked block in myeloid lineage progression, where *Brd9*-KO cells accumulated at the myeloid progenitor (GMP) stage (Fig 2g). Therefore, although both cBAF and ncBAF complexes are important for myeloid development, our data highlights their regulation of different stages of myeloid lineage progression.

However, in contrast to the diversity demonstrated within COMPASS and BAF subcomplexes, disruption of NuRD members (*Chd4, Mbd2*), NuRD associated Repressors (*Rbbp4, Hdac1*), Ncor (*Hdac3*) and heterochromatin regulators (*Setdb1* and *Atf7ip*) produced very similar phenotypes, characterised by accelerated granulocytic trajectories at the expense of erythroid and B-cell lineages (Fig 2f-g, j). Overall, these data suggest that a number of epigenetic repressors act to safeguard diversity of progenitor identities^24,25^, by limiting extensive myelopoiesis, thus ensuring balanced lineage output. The only exception to this behaviour was for *Rcor1*-KO, which, in line to its role in *ex vivo* bulk screens, generated erythroid lineage skewing and stalled differentiation trajectories with a marked accumulation of aberrant erythroid progenitors (Fig 2j). Finally, the Cohesin member *Stag2* and the arginine methyl-transferase *Prmt5* behaved similarly *in vivo*, with *Stag2*-KO blocking differentiation and *Prmt5*-KO reducing the numbers of progenitor cells (Fig S4d). Of note, many of the observed CF dependencies, especially for COMPASS and BAF complexes, do not relate to the expression patterns of these CFs suggesting that such lineage specificities are related to post-transcriptional regulation or to different regulatory properties or molecular requirements for CF activities within the different lineages (Fig S5).

### CFs regulate lineage specific expression programmes via TF chromatin accessibility

Next, to explore the molecular mechanisms underlying the marked CF lineage dependencies, we examined lineage specific expression programmes, including lineage-determining TFs and cell-type defining markers (Figs 3a, S6a). Depletion of the pro-myeloid complexes MLL4, cBAF and SET1 demonstrated a marked downregulation of myeloid programmes, perfectly recapitulating their lineage dependencies. In contrast, perturbation of the defining ncBAF subunit *Brd9* showed a mild disruption of the myeloid programme, a strong effect on key B-cell regulators and a marked upregulation of key progenitor genes, including the *Hoxa7-9* and *Meis1* transcription factors, that reflects the stalled myeloid progression observed for *Brd9*-KO. A similar upregulation of progenitor signatures was found in *Rcor1*- and *Wdr82*-KOs but this was combined with, respectively, upregulated erythroid and megakaryocytic TFs, which explains the accumulation of erythroid or megakaryocytic progenitors observed for *Rcor1*- or *Wdr82*-KO cells, respectively (Fig 2f, i-j). Finally, with the exception of *Hdac3*-KO, disruption of Repressors (*Setdb1, Atf7ip*) and NuRD components (*Rbbp4, Mbd2*) did not upregulate central myeloid regulators, suggesting that subtler but cumulative gene expression changes cause the observed myeloid bias upon their disruption. In keeping with this, GSEA analysis (Fig S6b) highlighted upregulation of inflammatory processes like “Interferon and JAK/STAT signalling” and, “Augmented activity of AP-1 transcription factors”, pathways known to enhance myeloid lineage outputs under inflammatory stimuli.

**Figure 3.**
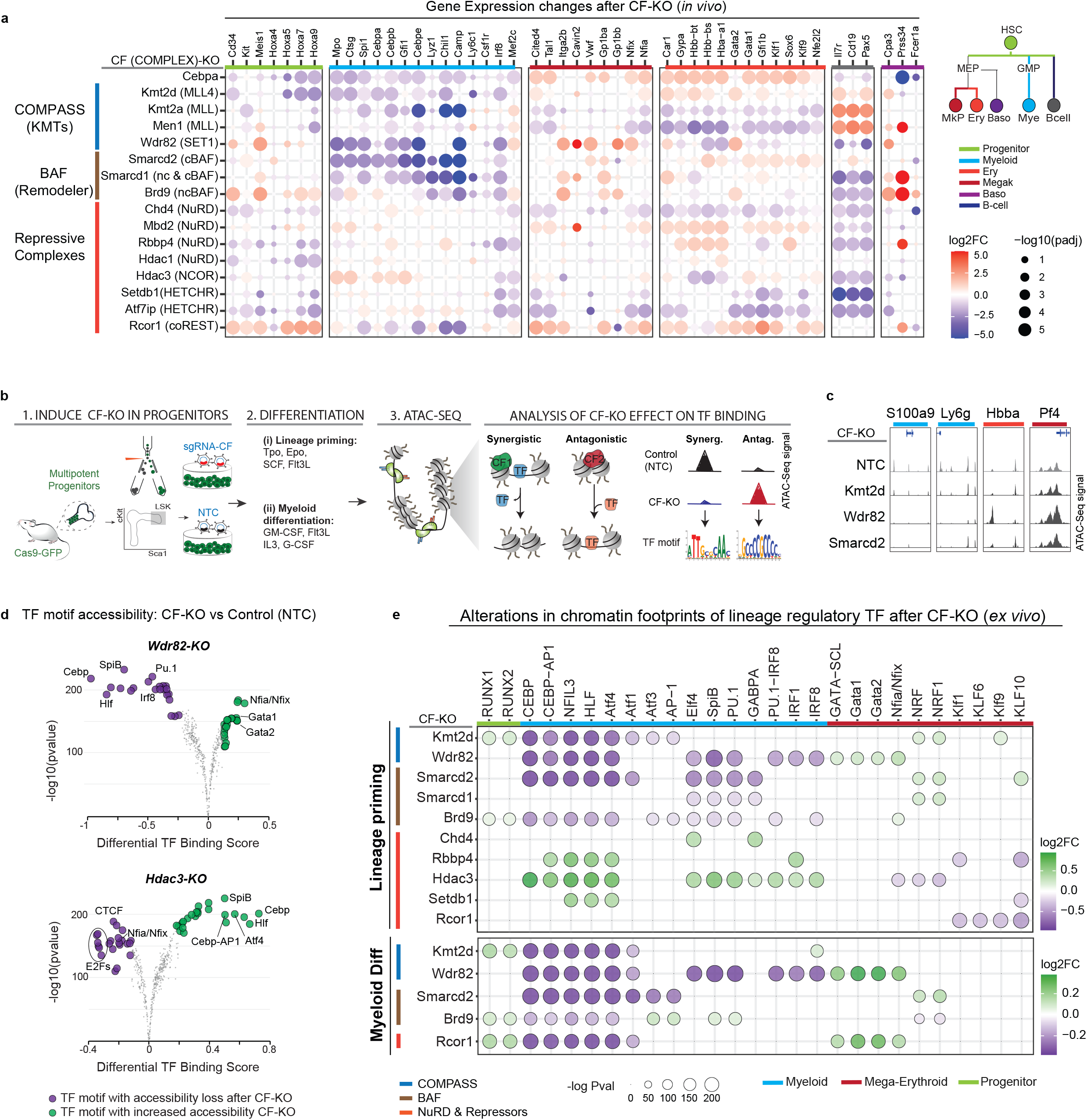
CFs control lineage specific expression programmes via regulation of TF chromatin accessibility. **(a)** Analysis of the effect of CF disruption (CF-KOs) on lineage specific expression patterns, comprising markers and transcription factors specific for Progenitor (Green), Myeloid (Light Blue), Megakaryocytic (Brown), Erythroid (Red), B-cell (Dark Blue) and Basophil (Purple) lineages. Dot colour and size relate to log2 fold change and -log10 adjusted p-value, respectively. **(b-d)** Evaluation of TF-CF dependencies **(b)** Schematic drawing outlining the experimental workflow: 1- CRISPR perturbation of CFs in multipotent progenitors; 2- Differentiation under lineage priming or myeloid conditions; 3- Detection of altered TF footprints with ATAC-seq. **(c)** Chromatin accessibility (ATAC-seq signal) for representative myeloid and mega-erythroid loci after disruption of specific CFs in *ex vivo* grown haematopoietic progenitors **(d)** Volcano plot showing differential TF footprinting analysis in Wdr82 and Hdac3 knockouts under lineage priming conditions. Analysis was performed with TOBIAS^26^ using NTC cells as baseline. Gained and lost accessibility for CF-KOs compared to NTCs are shown in green and purple, respectively. (**e**) Heatmap summarizing the effect of 10 CF-KOs on the motif footprints of TFs involved in lineage specification. Transcription factor motifs with increased accessibility (gain binding) upon CF-KO are shown in green and motifs with decreased accessibility (binding loss) are shown in purple as in (d). Dot colour and size relate to log2 fold change and -log10 adjusted p-value.

Intrigued by these findings, we decided to elucidate the TF-CF interactions that may explain the strong and lineage-specific CF dependencies observed *ex vivo* and *in vivo*. We therefore (1) used CRISPR to induce loss-of-function for 10 CFs, comprising COMPASS, BAF and Repressor complexes, in multipotent progenitors; (2) stimulated them with lineage instructive cytokines supporting lineage priming or myeloid differentiation and (3) utilised ATAC-seq to determine alterations in chromatin accessibility and, via motif and digital footprinting analysis^21^, to measure TF binding and activity, comparing CF-KO cells to NTC (Figs 3b-e, S7). This analysis revealed both synergistic and antagonistic connections between CFs and specific lineage-determining TFs. Myeloid-dependent CFs belonging to the COMPASS and BAF complexes regulated the accessibility of key myeloid TFs, however with different degrees of specificity. While MLL4 (*Kmt2d*-KO) is connected to Cebp, AP-1 and HLF myeloid TFs (all bZiP factors) cBAF (*Smarcd2*-KO) and SET1-COMPASS (*Wdr82*-KO) govern the accessibility of a larger repertoire of myeloid TF including Cebps, ETS (Pu.1) and IRF factors. Disruption of ncBAF (*Brd9*-KO) demonstrated a similar patter to cBAF, however the loss in accessibility was milder. Finally, in line with GSEA analysis of the *in vivo* KO phenotypes, disruption of NuRD (*Rbbp4*-KO) and other Repressors (*Hdac3*-KO and *Setdb1-*KO) induced increased accessibility of pro-myeloid TFs such as Cebp, HLF^27^ and AP-1^28^, suggesting that these repressive complexes block or attenuate a myeloid regulatory TF programme triggered by inflammatory cytokines (Fig 3e). Taken together these results highlight functional interactions between Chromatin Complexes and lineage determining TFs and suggest that the identity of the CF interaction-partners, at least in part, determines CF-specific lineage dependencies.

### Disruption of ncBAF leads to a pre-leukaemic accumulation of myeloid progenitors

Digital chromatin footprinting of CF-KOs highlighted TF-CF interactions determining the lineage dependences observed for specific CFs. However, this analysis did not explain the differential roles observed for cBAF vs ncBAF-complexes in myelopoiesis, where cBAF is required from an earlier myeloid priming stage and ncBAF is more critical for the progression of progenitors towards mature myeloid cells. Therefore, we speculated that this may have reflected incomplete recapitulation of myeloid lineage progression in our *ex vivo* ATAC-Seq system. To address this, we documented the dynamic binding patterns of ncBAF and cBAF along myeloid differentiation, performing ChIP-seq from immediately *in vivo* harvested early myeloid progenitors (GMPs) and mature myeloid cells (monocytes) (Fig 4a). In addition, as external comparators, we analysed Kmt2d- and Kmt2a-binding and included Erythroid and B-cell lineages (Fig S8a-c). Of note, since ChIP-grade antibodies for Smarcd2 weren’t available we instead assessed binding of Smarcb1, another defining subunit of cBAF. Differential peak binding analysis showed highly specific binding patterns for all CFs across the different lineages and differentiation stages (Figs 4b, S8c). Then, using motif analysis we analysed TF motifs associated with CFs in each differentiation-stage highlighting TF-CF interactions with lineage-instructing potential. This revealed a reshaping of the TF interaction landscape for the ncBAF complex during myeloid development, where Brd9 utilises broad TF partnerships (Cebp, Pu.1-Irf8 and Runx motifs) in myeloid progenitors, and a Cebp/Ap1 centric partnership in mature myeloid cells (Figs 4c, S8d). This suggests that the myeloid maturation defect observed in *Brd9*-KO cells may relate to impaired recruitment and activity of Cebp and Ap1 TF.

**Figure 4.**
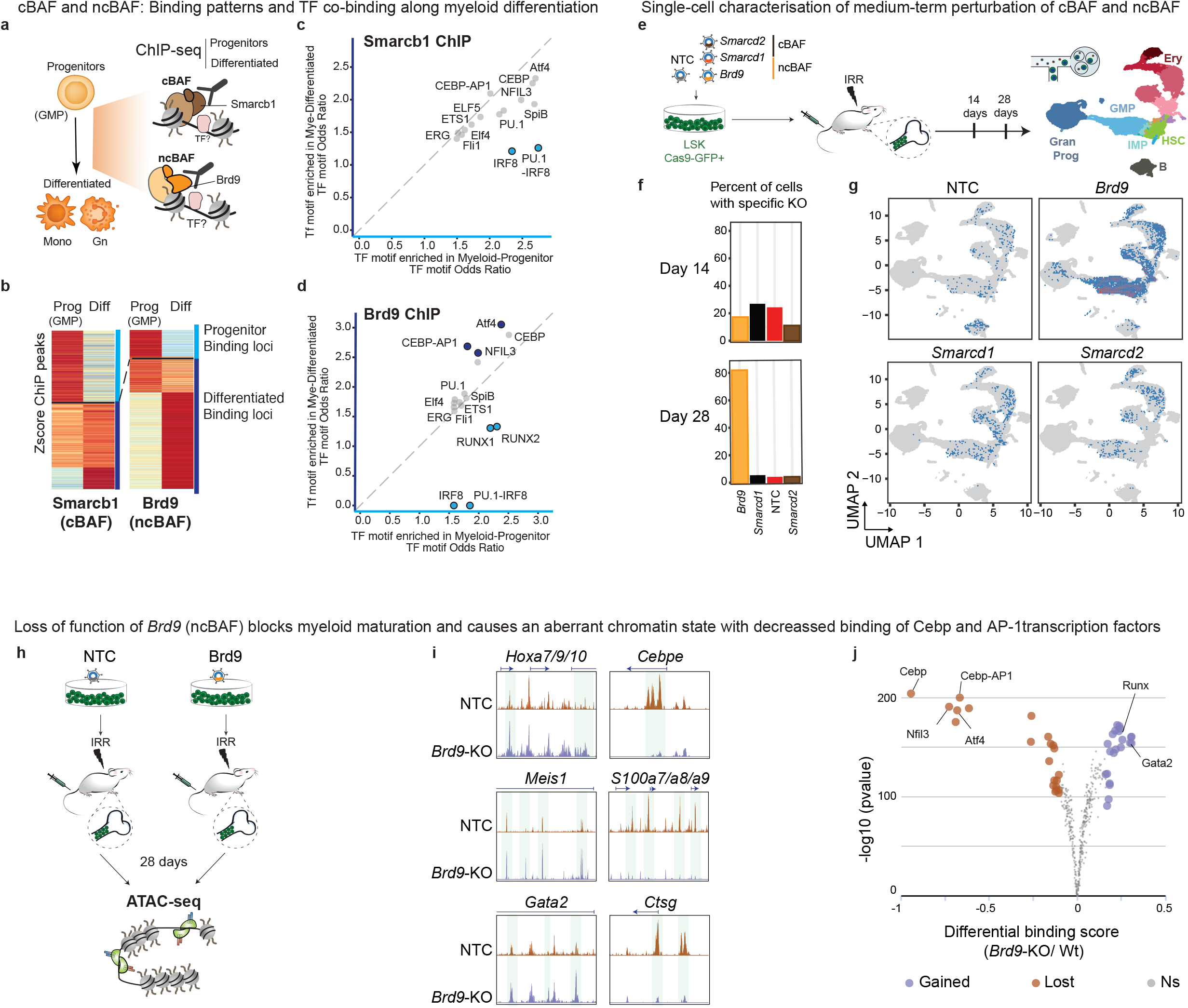
Disruption of ncBAF leads to a pre-leukaemic accumulation of myeloid progenitors. (a-d) Analysis of BAF targets using ChiP-seq. (**a**) Graphical drawing of the populations and BAF complexes interrogated with ChiP-seq. (**b**) Heatmaps showing specific binding of Smarcb1 (cBAF) and Brd9 (ncBAF) in early myeloid progenitors (GMPs: Lin-, ckit+, FcgR-III+, CD34+) and mature myeloid cells (bone-marrow monocytes: Lin-, Cd11b+). The heatmap shows the z-scores values for each peak, which are grouped using k-means clustering by their dynamic behaviour across different hematopoietic populations (see Fig S7c). **(c-d)** Scatter plots comparing the TF motif co-occurrence at early myeloid progenitor (GMP) versus mature myeloid specific loci (derived from 4b) for cBAF **(c)** and ncBAF **(d)**. The axis shows the Odds Ratio value for each TF motif. Light-blue coloured dots represent TF motifs specific for progenitors; dark-blue dots represents TF motifs specific for mature cells. All plotted TF motifs have adjusted p-values lower than 0.001 (e-g) Analysis of medium-term perturbations of BAF complexes using *in vivo* Perturb-seq. **(e)** Schematic drawing of the experiment competitively perturbing cBAF and ncBAF, with NTC controls, within the same transplant. **(f)** Barplots demonstrating numbers of cells derived from transplanted progenitors with a specific CF-KO at 14- and 28- days post-transplant. (**g)** UMAP showing the distribution of cells derived from NTC, *Brd9, Smarcd1* and *Smarcd2* perturbed progenitors at 28 days post-transplant. Cells are aggregated and the colour of each area represents the number of cells. The distribution of NTCs at day 14 post-transplant is shown as background in grey. (h-j) Analysis of Brd9 effects on chromatin *in vivo* using ATAC-seq. **(h)** Schematic drawing of the experiment. NTC and *Brd9*-KO cells were harvested at d28 for ATAC-Seq. **(i)** Genome browser tracks showing chromatin accessibility signal (ATAC-seq) at representative progenitor (*Hoxa* and *c-Kit* loci, left) and differentiated myeloid (*Cebpe, S100a7/a8/a9* and *Ctsg*, right) loci. **(j)** Volcano plot showing TF motifs with altered chromatin accessibility between NTC and *Brd9*-KO GMPs isolated at day 28 post-transplant. Gained and lost accessibility for TF are shown in blue and brown, respectively.

To better characterise the dynamics of cBAF and ncBAF requirement along myeloid development *in vivo*, we transplanted multipotent progenitors (LSK) obtained from Cas9 bone marrow transduced with a small CRISPR library containing control (NTC) *Smarcd2, Smarcd1* and *Brd9* sgRNAs and examined their differentiation patterns at 4 weeks with Perturb-seq (Fig 4c-e). This revealed a massive expansion of *Brd9-*KO clones, which outcompeted *Smarcd1-, Smarcd-*KO and NTC harbouring cells and mapped predominantly to the myeloid progenitor (GMP) compartment, confirming that Brd9 is required for myeloid progression and demonstrating that its disruption confers a competitive proliferation advantage. In consonance with this, chromatin accessibility profiling of *Brd9*-KO GMPs at 4 weeks post-transplant, in comparison with control GMPs, revealed an aberrant chromatin landscape, characterised by a decreased accessibility at loci associated with myeloid maturation including *Cebpe, S100a7-9* and *Ctsg* and increased accessibility at key progenitor loci such as *Hoxa7-9-10, Gata2* and *Meis1* (Fig 4h-i), genes whose expression is also upregulated in *Brd9*-KO progenitors (Fig 3a). In keeping with Brd9-associated TF motifs (derived from ChIP-seq – Fig 4d), comparison of digital footprinting between control and *Brd9*-KO GMPs revealed a marked loss of Cebp and Cebp-AP1 binding and increased activity of the progenitor-associated TFs Gata2 and Runx1 (Fig 4j). Together, these results demonstrate that Brd9 functionally interacts with Cebp and AP-1 TFs to drive later myeloid differentiation and suggests that *Brd9* depletion induces a preleukaemic state related, at least in part, to the failure to transition from TF programmes that maintain the progenitor state to later differentiation programmes.

### cBAF, COMPASS and Repressive CF complexes maintain leukaemic fitness through enforcing differentiation blockade in AML

Having suggested a tumour-suppressor role for ncBAF during myeloid development, we decided to explore any roles for key CFs in maintaining the aberrant chromatin landscapes that sustain established leukaemia cells. We therefore chose an aggressive *Npm1c*/*Flt3*-*ITD* model, driven by the two most common co-occurring mutations in AML that synergise to generate a highly corrupted chromatin landscape that recapitulates many aspects of human AML with the same genotype^29^. To interrogate CF function in this model, we isolated primary leukaemia cells from *Npm1c*/*Flt3-ITD/Cas9* mice, cultured them and used CRISPR to disrupt both cBAF and ncBAF members, the three COMPASS complexes and Repressive factors including NuRD, Ncor and heterochromatin regulators. The effects of such CF-KOs were then read-out by examining their growth dynamics and characterising their cellular states with Perturb-seq (Fig 5a). Remarkably, the majority of these CF-KOs led to fitness loss, reflected by reduced growth dynamics (Fig 5b). This was especially pronounced for cBAF and COMPASS members, revealing *Npm1c*/*Flt3-ITD* leukaemia to be highly dependent on the epigenetic activities regulated by these complexes. However, unlike previous reports for *MLL*-driven leukaemias^30,31^, our *Npm1c*/*Flt3-ITD* model did not show vulnerability to *Brd9* (ncBAF) disruption, highlighting that different leukaemic mutations produce specific chromatin states that are variably dependent on specific CFs. Analysis of the single cell expression patterns of the *Npm1c/Flt3-ITD* leukaemia identified cellular populations resembling granulocytic, erythroid, basophilic and megakaryocytic states (Fig 5c-d), that demonstrated decreased fitness when isolated and assessed (Figs 5e, S9a-c). Analysis of the single cell patterns generated by the disruption of dependent CFs in leukaemia provided a molecular explanation for the lower fitness observed for these CF-Kos (Fig 5f). Specifically, the depletion of MLL1-COMPASS (*Kmt2a*-KO and *Men1*-KO), NCOR/*Hdac3*, and Heterochromatin regulators *Setdb1* and *Atf7ip* induced transition towards the Granulocytic state. In contrast, loss-of-function of cBAF, MLL4-COMPAS and of SET1-COMPASS chromatin complexes pushed the leukaemia towards the basophil, erythroid and megakaryocytic states. Extending our analysis to a larger CF repertoire in search for novel therapeutic targets, we screened the requirements of more factors, for a total number of 50 CFs (Fig S9d), selected as having relevant roles in normal haematopoiesis. This uncovered a further number of vulnerabilities, including *Prmt1* and *Prmt5* as novel *Npm1c/Flt3ITD* leukaemia dependencies (Fig S9e), that may be amenable to therapeutic exploitation. These collective findings demonstrate that latent differentiation pathways exist in leukaemia which, if engaged, might facilitate replicative exhaustion (Fig 5c, f-g). Moreover, mechanistically this shows that leukaemic states co-opt CF normally critical for homeostatic differentiation to actively block differentiation, thereby preserving their replicative fitness.

**Figure 5.**
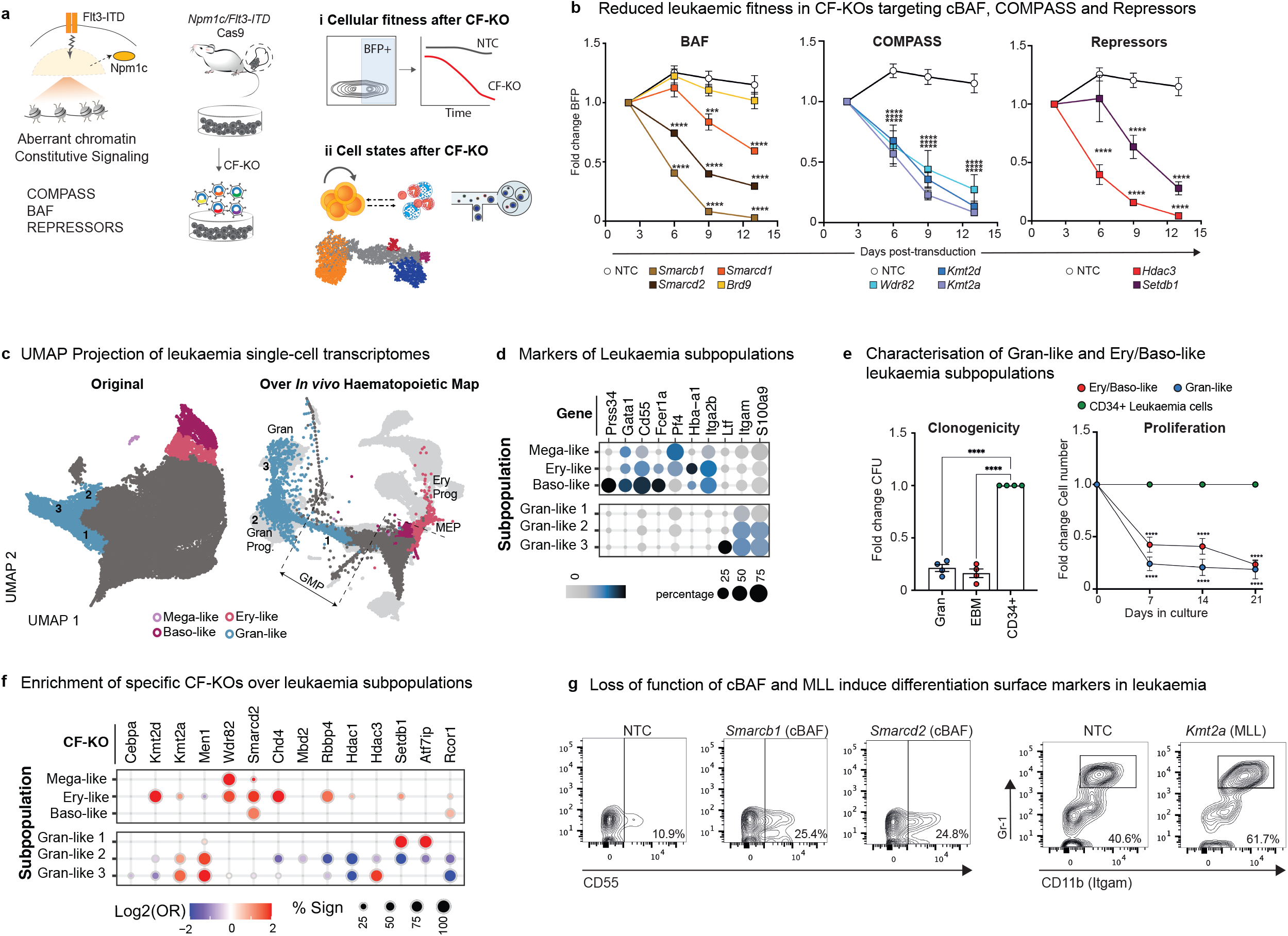
Canonical BAF, COMPASS and several chromatin Repressive complexes maintain leukaemic fitness through enforcing differentiation blockade in AML. (**a)** Conceptual drawing of *Npm1c/Flt3-ITD* corrupted chromatin and functionally and transcriptional methodologies to interrogate CF roles **(b)** Growth curves for leukaemia cells with specific CF-KOs. The cells expressing each sgRNA harbour a BFP reporter and the assay measures the change in the proportion of BFP expressing cells over time. **(c)** UMAP projection of single-cell transcriptomes from *Npm1c/Flt3-ITD* primary leukaemia. Colour-coded clusters correspond to cells with differentiation-like signatures: Granulocytic-like (blue), Erythroid-like (red), Basophil-like (purple) and Megakaryocytic-like (violet). **(d)** Expression of differentiation makers in the leukaemia subpopulations. **(e)** Clonogenic and proliferation assays for the various leukaemia subpopulations. These were isolated following the strategy in panel S9c. **(f)** CF-KOs induce differentiation trajectories in leukaemia towards: Granulocyte-like, Basophil-like, Erythroid-like and Megakaryocytic-like subpopulations. Enrichment analysis was performed for each CF-KOs across the leukaemic clusters. Dot colour and size relate to the log2 odds ratio and the percent of significant enrichments versus NTCs, respectively. **(g)** Disruption of cBAF (*Smarcb1*-KO and *Smarcd2*-KO) and MLL (*Kmt2a*-KO) induce expression of surface differentiation markers in *Npm1c/Flt3-ITD* leukaemia.

### CF enforce differentiation blockade in AML through novel TF interactions

Finally, to interrogate the molecular mechanisms that underpin the requirement for the cBAF and MLL-COMPASS complexes in *Npm1c*/*Flt3-ITD* AML, and to explain their differing role from normal haematopoiesis, we compared the genome-wide binding patterns of *Smarcb1* (as an exemplar of cBAF), *Kmt2a* (COMPASS-MLL) and *Kmt2d* (COMPASS-MLL4) by ChIP-Seq across the leukaemia, normal myeloid progenitor (GMP) and *in vivo* and *ex vivo*-derived mature myeloid subsets (Fig 6a-c,e). Of note, these analyses demonstrated a marked redistribution of the cBAF and MLL1/4-COMPASS complexes upon leukaemia induction (Fig 6c, e). Binding patterns across the three cell types identified 3 groups; (1) Loci that are bound only in leukaemia (Leukaemia), (2) that are common to leukaemia and normal cells (Common) and (3) that are only bound in normal cells (Normal). Leukaemia-specific Loci were demonstrated to be enriched in relevant molecular functions, such as Tyrosine Kinase Signalling, related to the FLT3-ITD mutation (Fig S10). Moreover, motif analysis across the binding patterns in the three separate groups (Fig 6d) highlighted corrupted TF-CF interactions in leukaemia; where Stat5a scored as the top TF leukaemia-specific interactor for MLL1/4 and cBAF complexes. By contrast, Pu.1 and IRF factors were associated with cBAF (Smarcb1 binding) specifically in normal cells and, MLL4 shared Cebp, Nfil3 and Hlf interactions in both normal and leukaemia scenarios (Fig 6d, f).

**Figure 6.**
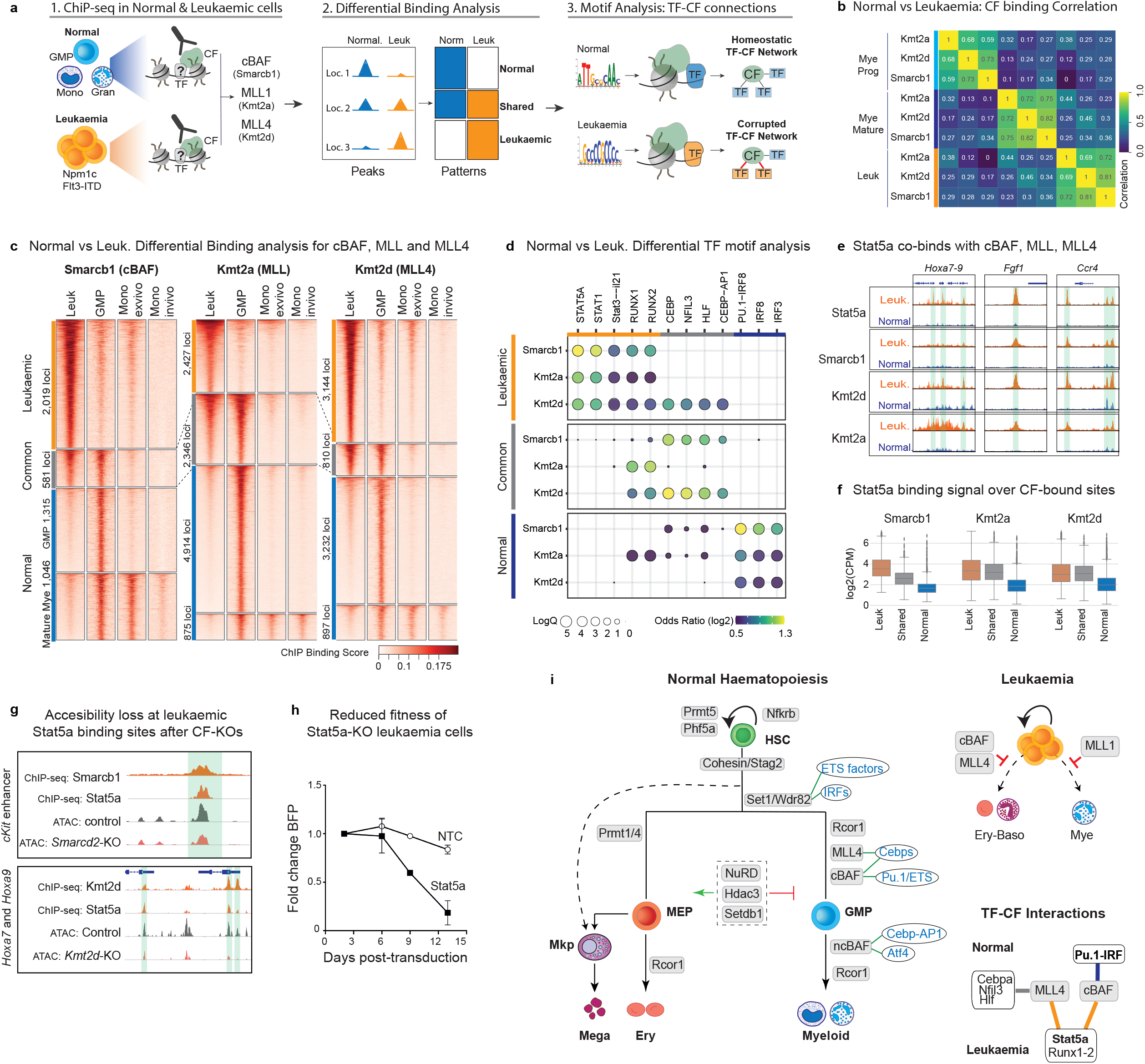
Functional analysis of CFs in *Npm1c/Flt3-ITD* AML identifies CF-dependencies and leukaemia specific TF-CF interactions. **(a)** Conceptual drawing showing ChIP-seq based strategy to identify corrupted TF-CF interactions in leukaemia **(b)** Correlation heatmap of ChIP-seq signals of cBAF (Smarcb1), MLL (Kmt2a) and MLL4 (Kmt2d) chromatin complexes in normal myeloid cells (Progenitors and Mature) and *Npm1c/Flt3-ITD* leukaemia cells. **(c)** Heatmap**s** highlighting alterations between the leukaemia and normal binding patterns for cBAF (Smarcb1), MLL (Kmt2a) and MLL4 (Kmt2d). The assayed populations are primary *Npm1c/Flt3-ITD* leukaemia cells, *in vivo* myeloid progenitors (GMPs), *in vivo* monocytes and *ex vivo*-derived primary monocytes. **(d)** Motif Enrichment analysis of cBAF (Smarcb1), MLL (Kmt2a) and MLL4 (Kmt2d) binding patterns specific for leukaemia (Leukaemia), common between leukaemia and myeloid progenitors (Common), and specific for normal myeloid cells (Normal - including both progenitors and mature cells). **(e-f)** Stat5a co-binds with cBAF, MLL and MLL4 at leukaemia specific loci: **(e)** Genome-browser snapshots at known leukaemia loci **(f)** Boxplot showing Stat5a binding signal at leukaemic, common and normal loci (defined in figure 6c) for cBAF, MLL and MLL4. **(g)** Disruption of cBAF and MLL4 reduces the chromatin accessibility of Stat5a binding sites at leukaemia specific loci. **(h)** Loss of Stat5a compromises the proliferative fitness of *Npm1c/Flt3-ITD* leukaemia cells. Growth curves for leukaemia cells with *Stat5a*-KO and control (NTC). The cells expressing each sgRNA harbour a BFP reporter and, the assay measures the change in the proportion of BFP expressing cells over time. (i) Conceptual summary of the main findings: (left) Roles of CF in lineage trajectories during haematopoiesis, with TF interactions demonstrated by this study highlighted in blue. (top right) Corrupted roles of CF in leukaemia. Note the switch in roles from normal haematopoiesis where CF generally facilitate differentiation, to leukaemia, where they actively maintain differentiation blockade. (bottom right) Switch in TF-CF interactions from normal to leukaemia highlighted in lower right panel.

ChIP-seq of Stat5a confirmed the motif analysis and demonstrated an increased binding of Stat5a at Leukaemia loci, including binding at known key pro-leukaemia genes such as *Hoxa7* and *Hoxa9* (Fig 6d). Moreover, chromatin accessibility profiling of leukaemia cells disrupted for cBAF (*Smarcd2*-KO) or COMPASS-MLL4 (*Kmt2d*-KO) confirmed the requirement for cBAF and MLL4 to maintain optimal accessibility at critical Stat5a bound loci (Fig 6g). Finally, loss of function of *Stat5a* in *Npm1c/Flt3-ITD* cells significantly reduced the proliferative fitness of this AML, confirming the importance of these TF-CF switches for AML maintenance (Fig 6i). Taken together these findings demonstrate how individual CF required for lineage determination in normal myeloid differentiation engage in corrupted interactions with alternative TF partners to promote differentiation blockade, thereby maintaining cellular fitness in AML.

## Discussion

Chromatin factors mediate a wide repertoire of epigenetic activities with key instructive genome regulatory roles. However, unlike TF, whose expression patterns often mirror their functional activity, CF have a more stable, uniform expression pattern, suggesting a more generic role in gene regulation that is deployed to cell-type specific loci via lineage-determining TFs. Here, combining bulk and single cell perturbations we uncover diverse instructive roles, across multiple haematopoietic lineages, for specific CFs and CF-complexes, with CF even able to phenocopy (*Kmt2d*) the loss-of-function of lineage-determining TFs, such as the master-myeloid regulators *Cebpa* and *Pu*.*1*. Further demonstrating the complex nature of this CF specificity, related chromatin complexes that mediate similar epigenetic activities, including COMPASS H3K4-methyltransferases or different BAF-complex configurations, demonstrate remarkably different roles. This shows that, despite depositing similar epigenetic marks, the different COMPASS KMTs are not redundant and suggests that they individually regulate specific H3K4me patterns, or that their distinct roles are mediated via catalytic-independent activities, as has been reported for other CFs^32^. In stark contrast, apparently unrelated repressive CF belonging to the NuRD, Ncor and Heterochromatin complexes could be shown to have uniform functional activities with all attenuating excessive granulopoiesis, likely by dampening chromatin responses to inflammatory mediators.

These observations pose the question of what then underlies this CF specificity? Unlike TFs, CFs lack specific DNA binding domains, however many proteins and complexes contain reader domains that bind loci with specific epigenetic configurations and some central CFs like BAF remodelers or COMPASS methyl-transferases have been shown to interact physically or functionally with specific TFs^33,34,35,36,37^. Here, we have functionally mapped TF-CF connections for 10 CF with strong lineage dependencies. These confirm that specific CFs are required to mediate the binding of key lineage-determining TFs in particular cellular contexts and highlight specific complexes, including BAF or SET and MLL4 COMPASS, as central regulators of lineage specific chromatin accessibility patterns. Of note, neither SET1 nor MLL4 possess chromatin remodelling activity, therefore their effects on chromatin accessibility must, in-turn, be mediated by the recruitment of chromatin remodelers; either directly via direct protein-interaction or indirectly through configuring a local pattern of histone methylation that recruit remodelers via their own reader modules^38^. Supporting this, we observed strong phenocopy between MLL4 and canonical BAF at both the cellular and molecular level, with both complexes synergising to induce myeloid regulatory programmes, a pattern reported in other systems^39^. Interestingly, our identified TF-CF connections show different degrees of specificity, with factors like SET1 connected to a large TF repertoire and factors like MLL4 having a selective partnership centred around bZIP TFs including Cebp or Nfil3. In addition, as another regulatory layer, our demonstration that chromatin repressors block the accessibility of pro-myeloid TFs activated in our cytokine-rich experimental systems, suggests these CF to buffer excessive extracellular stimuli, thereby ensuring a balance of haematopoietic cell production.

Studying myelopoiesis *in vivo*, we could also demonstrate the potentially dynamic nature of these CF-TF interactions; for example, the alteration of TF-interactions for the ncBAF subunit Brd9 that, upon myeloid maturation, changes from a broad TF repertoire to specific Cebp and Ap1-factors. Furthermore, we could functionally validate these interactions through perturbation of ncBAF, which blocked myeloid maturation through disabling the activity Cebp, AP-1 and Atf4 factors known to be involved in myeloid maturation^40,41^. Of interest, this perturbation led to the accumulation of myeloid progenitors with a preleukaemic gene-expression programme, and in this respect *Brd9*-loss may mimic the effects of *CEBPA* mutations, that occur both somatically and in the germline, where they generate either a genetic predisposition to AML development or an initiating mutation^42,43^. Furthermore, aberrant splicing of *BRD9*, to include a so-called “poison exon” leading to its degradation, has been previously described in the AML-precursor lesion, MDS^44^.

Studying CF requirement in AML through a combination of functional, CiTE-seq and Perturb-seq experiments we have been able to identify known (MEN1, PRMT5^45,46^) and novel leukaemic dependencies, that may be amenable to inhibition/targeting, including Prmt1, Hdac3, Setdb1 and Kmt2d. Mechanistically, our data also highlight opposing functions for the same CF in normal and malignant haematopoiesis. CFs such as COMPASS members and cBAF remodelers are critical for the balanced differentiation of specific lineages, however the same CF are critical for the maintenance of leukaemia fitness, where they act predominantly to maintain differentiation blockade (Fig 5i). Moreover, using comparative chromatin profiling we show that the proleukaemic roles of COMPASS and BAF complexes are mediated by the rewiring of their TF interactions, where these factors decrease interaction with IRF and PU.1 TF networks and gain novel connections to Stat5a and Runx factors, proving the relevance of these interactions by showing loss of leukaemia fitness upon *Stat5a*-KO. These observations also have clinical implications; TF and CF have pleiotropic effects across multiple tissues, but targeting specific TF-CF interactions will likely have much lower toxicity and higher specificity. However, designing such therapies will require a detailed knowledge of the specific structures and mechanisms governing individual TF-CF and these studies are warranted^47^.

Finally, our combination of functional, large-scale CRISPR-screening with downstream single cell analysis *in vivo* could be readily deployed to assess the role of other general classes of proteins across normal and malignant haematopoiesis. Moreover, it could also be adapted to perform similar physiological screens in other organ and tumour systems.

Taken all together, our studies show that Chromatin Factors are a highly dynamic and specific regulatory layer that should be on an equal weighting with TFs when studying cell fate decisions in both normal and malignant haematopoiesis, and we expect in other systems. Our studies lay the basis for additional, in-depth interrogation of specific CF-TF functions, using multidisciplinary approaches ranging from *in vivo* functional approaches to protein-protein interactions, that we feel are warranted to further elucidate their function.

## Figure legends

**Supplementary Figure 1** | **Characterisation of CRISPR Screen systems**. (**a**) Comparison between the expression profiles of the different readout populations from our screens and lineage specific signatures from 3 different studies, named along the bottom of the graph. We performed enrichment analysis between the screen signatures and the reference signatures. Dot colour and size relate to log2 odds ratio and -log10 adjusted p-value (key to right of panel), respectively. CLP – Common Lymphoid Progenitor, MkP – Megakaryocyte Progenitor. (**b**) Comparison between the expression profiles of the different readout populations and a single-cell map of normal haematopoiesis from Izzo *et al*^48^. Bulk transcriptomic signatures from our *ex vivo* systems, derived from FACS-sorted populations, were projected on the single-cell map from Izzo *et al*. **(c)** Example distribution of the CF lineage scores calculated from the Cas9 (green border) and Non-Cas9 (grey) populations. **(d)** Results from 2 CRISPR screen replicates for selected CFs. **(e)** Comparison of CF dependencies between Myeloid differentiation from LSK progenitors (y-axis) and Terminal Myeloid Differentiation from GMPs (x-axis), similar to Fig 1c. For each gene, lineage scores are shown aggregated across all guides and libraries. Genes with significant changes are shown with a red cross. Genes that did reach significance thresholds are shown as black dots, aggregated NTCs are shown with a blue cross. Data for non-Cas9 cells are shown in the background using a 2-dimensional yellow-blue density map. **(f)** Lineage scores for all hits. The colour of each dot represents the aggregated lineage score. The size represents the number of significant guides, as per key to the right.

**Supplementary Figure 2** | **Validation of the effects of individual CF-KOs. (a-b)** Heatmap showing changes in representative populations for each CF-KO compared to Non-Targeting Control (NTC) under lineage-priming **(a)** and myeloid differentiation and terminal myeloid maturation **(b)**. Gates and values for the selected populations are derived from figures S2c-d. **(c-d)** FACS plots showing validation results for individual CF-KOs under lineage priming conditions **(c)** and Terminal Myeloid Differentiation **(d)**. These validations were performed in different batches. Each batch included a Non-Targeting Control condition. The results were always normalised to the NTC included in each batch.

**Supplementary Figure 3** | **Characterisation of the *in vivo* and *ex vivo* systems based on single-cell transcriptomics**. (**a**) Comparison of expression profiles from our *in vivo* single-cell clusters and external cell-type signatures from^48^. **(b)** Number of cells per cell type. **(c)** Fraction of cells in each cluster harbouring Non-Targeting Guides 1 and 2 (NTC1 and NTC2) across the different Perturb-seq batches. **(d)** Number of cells with a specific CF-KO. **(e-f)** Single-cell characterization of the *ex vivo* Perturb-seq system. **(e)** UMAP projection of the *ex vivo* single-cell transcriptomes over the *in vivo* UMAP coordinates. **(f)** Comparison of expression profiles from our *ex vivo* single-cell clusters and external cell-type signatures from^48^.

**Supplementary Figure 4** | **Extended *In vivo* Perturb-seq analysis of Chromatin Regulatory Complexes during lineage specification**. (**a) (**Left) Enrichment analyses of CF-KOs across main cellular lineages *in vivo*. Myeloid represents aggregated population of IMPs, GMPs, Monocytes, Granulocyte Progenitors and Granulocytes. Erythroid comprises MEPs, Erythroid Progenitors and Mature Erythroid cells. (Right) Enrichment analyses of CF-KOs between early myeloid (IMPs, GMPs & Granulocyte progenitors) and early erythroid (MEPs and Erythroid Progenitors) branches *in vivo*. OR = Odds ratio. %sign = percent of significant enrichments versus NTCs. The colour of each dot represents the log2(OR), the size represents the percent significant cells, as per key to the bottom right. (**b**) Enrichment analyses of CF-KOs across main cellular lineages *ex vivo*. OR = Odds ratio. %sign = percent of significant enrichments versus NTCs. Key as before. **(c)** Heatmap summarising trajectory analysis for CF-KOs along myeloid and erythroid branches ordered from HSCs to mature lineage using pseudotime. Colours are given by signed negative log10 p-values (for p<0.01) generated by a t-test between targeting and non-targeting control populations such that negative values correspond to reduced differentiation capability and positive values correspond to increased differentiation capability. **(d)** Distribution of cells with specific CF-KOs from the *in vivo* system plotted on the UMAP generated from *in vivo* single-cell transcriptomes. The distribution of NTCs is shown as background in grey in all plots. Cells are aggregated and the colour of each area represents the density of cells in each area.

**Supplementary Figure 5**| **Expression profiles of selected Chromatin Factors in different haematopoietic lineages**. Barplots showing the expression levels of selected Chromatin Regulators. The bars represent the normalized read counts taken from a RNA-seq dataset^49^.

**Supplementary Figure 6** | **Extended Analysis of transcriptomic effects of CF-KOs. (a)** Analysis of the effect of Chromatin factor disruption (CF-KOs) on lineage specific expression patterns, comprising markers and transcription factors specific for Progenitor, Myeloid, Erythroid, Megakaryocytic, Basophil and B-cell lineages. The colour of each dot represents the log2 fold change, the size represents the –log10 adjusted p-value, as per key to the bottom right. **(b)** Gene set enrichment analysis (GSEA) of differentially expressed genes in knockouts of factors belonging to repressive complexes. The colour of each dot represents normalized enrichment score, the size represents the –log10 adjusted p-value.

**Supplementary Figure 7** | **Analysis of CF-KOs on chromatin and TF motif accessibility**. (top) MA plots demonstrating differential accessibility analysis between CF-KOs and Control (NTC). Up- and down-regulated genomic loci are indicated in red and blue, respectively. (bottom) Volcano Plots showing the differential footprinting analysis between CF-KOs and Control (NTC). Gained and lost footprints are indicated in red and blue, respectively.

**Supplementary Figure 8** | ***In vivo* binding patterns of BAF and COMPASS complexes. (a)** CF binding at representative loci in myeloid and erythroid progenitors **(b)** Distance of identified peaks to transcriptional start sites (TSS). **(c)** Heatmaps showing lineage specific binding patterns for each CF. **(d)** TF motif co-occurrence in lineage specific binding patterns of cBAF, ncBAF, MLL and MLL4. TF motifs are sorted by their Odds Ratio Values (y-axis) in Kmt2a- Kmt2d- Brd9- and Smarcb1- lineage specific peaks: GMP, Mye (GMP & Monocytes), MEP, Ery (MEP & Erythrocytes) and B-cells. The colour scale reflects the –log10 p-adjusted values for each TF motif.

**Supplementary Figure 9. Extended analysis of Chromatin Factor roles in Npm1c/Flt3-ITD Leukaemia. (a)** UMAP projection showing scaled CITE-seq signal for 9 surface markers in leukaemic cells. The UMAP is based on the projection of the leukaemic single-cell transcriptomes onto the *in vivo* data shown in Fig 2a. **(b)** Heatmap showing scaled CITE-seq signal across low fitness leukaemia clusters for 9 surface markers. **(c)** Example sorting strategy of leukeamic subpopulations showing traits of differentiation into Granulocytic (Gran) or a mixed Erythroid-Basophil populations. **(d)** Enrichment analyses of all CF-KOs across leukaemia subpopulations. Growth curves of *Prmt1*- and *Prmt5*-KO cells. The cells expressing each sgRNA harbour a BFP reporter and, the assay measures the change in the proportion of BFP expressing cells over time.

**Supplementary Figure 10. Functional enrichment of Smarcb1, Kmt2a and Kmt2d leukaemia specific binding loci**. Bar graphs of enriched terms across input gene lists, sorted by p-values. Functions with particularly high relevance for *Npm1c/Flt3-ITD* leukaemia are highlighted in bold. Targets connected to leukaemic specific peaks (nearest TSS) were run in Metascape^50^.

## EXPERIMENTAL PROCEDURES

### Mouse models

All animals were housed at Centre for Applied Medical Research (CIMA) and the Anne McLaren building at the University of Cambridge. All animal procedures were completed in accordance with the Guidelines of the Care and Use of Laboratory Animals and were approved by the Institutional Animal Care and Use Committees at University of Navarra, Spain, and the Animal Welfare Ethical Review Body at the University of Cambridge, UK. C57BL/6J (Jackson Laboratory #JAX_000664) and Gt(ROSA)26Sortm1.1(CAG-cas9*/EGFP)Rsky (Jackson Laboratory #JAX_026179) were used for all experimental procedures. The *Npm1c/Flt3-ITD/Cas9* model has been extensively described previously^29, 76, 77^.

### CRISPR Libraries Construction

sgRNA-CRISPR library targeting 550 chromatin regulators (Suppl Table 1) was ordered from IDT Technologies and cloned using Gibson assembly in CRISP-seq backbone (Addgene #85707). The Gibson assembly product was electroporated in Endura ElectroCompetent cells following the manufacturer’s protocol (Endura #60242-2) and plated on Bioassay LB agar plates. After 20 hours at 30 ºC bacteria were harvested and plasmid preparations were performed using Macherey-Nagel™ NucleoBond™ Xtra Midi EF (Macherey-Nagel **#**740420.50).

### Cloning of individual sgRNAs

Individual oligos were cloned in the CRISP-seq backbone (Suppl Table 1) following a Golden-Gate protocol by performing 20 cycles of digestion and ligation with Esp3I and T4 DNA ligase. Golden-Gate reactions were heat inactivated and transformed into NEB Stable Competent E. coli and grown on LB agar plates for 20 hours at 30 ºC. Two individual colonies were picked, grown in liquid LB + Ampicilin and processed with the ZymoPURE Plasmid MiniPrep Kit. The plasmid preps were Sanger sequenced using a U6 Forward (Fw) primer.

### Lentiviral production

HEK293T cells were seeded at 25 million cells in 175 cm^2^ tissue-culture treated flasks 16 hours prior to transfection and cultured in DMEM supplemented with 10% FBS and 1% Pen/Strep. The following day, media was replaced with DMEM + 10% FBS without Pen/strep and cells were transfected with the CRISP-seq plasmid backbones and 2^nd^ generation lentiviral packaging plasmids, pMD2-G (Addgene #12259) and psPAX2 (Addgene #12260) using lipofectamine 3000 following the manufacturer’s protocol. After 10 hours, media was replaced with Opti-MEM (ThermoFisherScientific #31985070) plus 1% Pen/Strep. The viral supernatant was collected within 48 hours, centrifuged at 300 g for 5 minutes to remove cell debris, filtered using 0.45 uM filters and concentrated with a Centricon with 100 kDa cutoff via centrifugation at 3000 xg at 4 ºC. Concentrated viral preps were aliquoted and stored at −80 ºC.

### Bulk CRISPR screens *ex vivo*

#### Isolation of murine hematopoietic progenitors (LSK) from bone marrow

Femora, tibiae, ilia, humerus, sternum and scapula were harvested from C57BL/6J and ROSAxCas9 mice, crushed with a pestle and mortar using cold (4 ºC) autoMACS Running Buffer and filtered through a 70 µM strainer. Red Blood Cells were lysed using RBC Lysis Buffer and cKit+ cells were enriched using magnetic beads, following the manufacturer’s protocol. The cKit enriched fraction was stained with BV510 anti-mouse Lineage (B220, CD3, CD11b, Gr1, Ter-119), APC anti-mouse CD117 and PE anti-mouse Sca1 (Suppl Table 2). Lin-/cKit+/Sca1+ cells (LSKs) were FACS-sorted in 1 mL of DMEM/F12 + 1X Pen/Strep, centrifuged at 350xG for 10 minutes, resuspended in culture media (see below) and seeded for *ex vivo* bulk screens or *in vivo* transplantation and Perturb-seq.

#### Ex vivo CRISPR screens

After FACS-sorting, multipotent (LSK) or myeloid (GMP) progenitors were resuspended at 250 cells/ul in DMEM/F12 plus 1% PSG, PVA 87%, ITS-X, 1X Hepes, 100 ng/mL murine Tpo and 10 ng/mL murine SCF and plated in 96w plates with 100 ul (25,000 cells) per well. Immediately after plating, cells were infected with the Lenti-CRISPR-BFP library previously titrated to reach a MOI of ∼20% infection. After 12 hours, two volumes of fresh medium were added. Then, at 48 hours post-infection, the cells were diluted in 2 mL of one of 4 (below) “Screen Media”, plated in 6-well plates and cultured for the specific time (see below):

1. <u>Stem cell vs Differentiation:</u>
  - Time: 9 days
  - Media. Complete DMEM/F12 (1% PSG (Gibco, PVA 87%, ITS-X, 1X Hepes) + 100 ng/mL murine Tpo +10 ng/mL murine SCF
  - FACS readout
2. <u>Lineage Priming:</u>
  - Time: 5 days
  - Media: Complete DMEM/F12 + 100 ng/mL murine Tpo + 1 ng/mL murine Flt3L + 10 ng/mL murine SCF
  - FACS Readout
3. <u>Myeloid differentiation:</u>
  - Time: 4 days
  - Media: IMDM, 20% FBS, 1% PSG, 10 ng/mL murine GM-CSF, 10 ng/mL murine SCF, 5 ng/mL murine G-CSF, 5 ng/mL murine IL-3, 5 ng/mL murine IL-6, 5 ng/mL murine IL-5, 5 ng/mL murine Flt3L, 2 ng/mL murine Tpo and 2 U/mL Epo
  - FACS Readout
4. <u>Terminal Myeloid Differentiation:</u>
  - Time: 2 days
  - Media: IMDM, 20% FBS, 1% PSG, 10 ng/mL murine GM-CSF, 10 ng/mL murine SCF, 5 ng/mL murine G-CSF, 5 ng/mL murine IL-3, 5 ng/mL murine IL-6, 5 ng/mL murine IL-5, 5 ng/mL murine Flt3L
  - FACS Readout

#### FACS Readouts

Cultures were harvested by centrifugation at 300 g for 5 minutes and washed twice with 1X cold PBS. Then the cell pellets were stained with the Readout specific cocktails (see below). Viable (TOPRO or Propidium Iodide), BFP+ cells (containing CRISPR guides) were gated from Cas9 (GFP+) and Non-Cas9 (GFP-) fractions and from each fraction the readout populations (see below) were sorted in 1.5 mL tubes containing PBS + 0.5% BSA.

1. <u>Stem cell vs Differentiation:</u>
  - Stain with BV510 lineage (B220, CD3, CD11b, Ly-6C/Ly-6G, Ter-119) APC anti CD117 and PE anti Sca1.
  - Sort: Multipotent Progenitor fraction (Lin-, cKit+, Sca1+) vs Differentiated fraction (Lin-, cKit+, Sca1-).
2. <u>Lineage Priming:</u>
  - Stain with BV510 lineage (B220, CD3, CD11b, Ly-6C/Ly-6G, Ter-119) APC anti CD117, PE anti Sca1 and PercPCy5.5 anti-mouse FcgRIII.
  - Sort: Myeloid Progenitor (Lin-, cKit+, Sca1-, FcgRIII+) vs Mega-erythroid progenitor (Lin-, cKit+, Sca1-, FcgRIII-).
3. <u>Myeloid Differentiation:</u>
  - Stain with PercPCy5.5 anti-mouse FcgRIII and PE-Cy7 anti CD11b.
  - Sort Myeloid cells (FcgRIII+, CD11b+) vs Non-myeloid fraction (FcgRIII-, CD11b-).
4. <u>Terminal Myeloid Differentiation</u>
  - Stain with BV510 anti Gr1 and PECy7 anti CD11b.
  - Sort Mature Myeloid fraction (CD11b+, Gr1+) vs Immature Myeloid fraction (CD11b-, Gr1-).

#### Bulk CRISPR Library Preparation

After cell sorting, cells were centrifuged at 700 g for 10 minutes with low brake, the supernatant was removed leaving ∼ 5 uL buffer cover and cell pellets were stored at −80 ºC until the library prep process was started.

For library prep: cell pellets were subjected to 3 cycles of freeze/thaw and lysed in 40 uL of lysis buffer (0.2% SDS and 2 uL of Proteinase K) at 42 ºC for 30 min. After the incubation, gDNA was isolated with 2X SPRI Cleanup. CRISPR sequencing libraries were prepared from purified gDNA with a 2-step PCR protocol:

- <u>1st PCR:</u> 200 ng gDNA 2.5 uL of 10 uM Read1-U6 and Read2-scaffold primers (Suppl Table 3), 25 uL of Kappa HIFI 2X and water to 50 uL total. PCR cycling conditions: 3 minutes at 98 ºC, followed by 10 s at 98 ºC, 10 s at 62 ºC, 25 s at 72 ºC, for 20 cycles; and a final 2 minutes extension at 72 ºC.
- <u>Purification</u>: 1X SPRI Cleanup. Elute in 25 uL.
- <u>2nd PCR:</u> 15 ng PCR1 product, 12.5 uL of Kappa HIFI 2X, 1.25 uL of each of the 10 uM P5-index and P7-index indexing primers (Suppl Table 3), and water to 25 uL total per reaction. PCR cycling conditions: 3 minutes at 98 ºC, followed by 10 s at 98 ºC, 10 s at 62ºC, 25 s at 72 ºC, for 10 cycles; and a final 2 minutes extension at 72 ºC.
- <u>Final purification</u>: 1X SPRI Cleanup. Elute in 25 uL.

Library QC was performed by measuring the library DNA concentration with Qubit dsDNA high sensitivity assay kit (range 1-5 ng/uL) and the library size was assessed by Tape Station, using the D1000 screen take (peak at bp)

Sequencing: 10M reads per sample in a NextSeq 1000 instrument with Rd1:50, i7:8; i5:8, Rd2:50

#### Computational analysis of FACS-based CRISPR screens

All analyses in this section were performed in R (version 4.0.2). Biological replicates were merged by summing the counts. Aggregated guide counts were normalised by first calculating normalising factors using the function *calcNormFactors* from edgeR (version 3.32.1)^51^, applied to the counts of only non-targeting guides in each aggregated sample. Counts were then transformed to counts per million (CPM) using the above calculated normalising factors, and further log_2_ normalised using the function *voom* from limma (version 3.46.0)^52^. A raw lineage score comparing pairs of populations (A and B) was then calculated by subtracting the log_2_ CPM of population A from the log_2_ CPM of population B, effectively resulting in a log_2_ fold change. This analysis was performed separately for each guide library. To assess the significance of population differences, we next calculated the probability of observing a given raw lineage score (log_2_ fold change) in the Cas9 data given the Non-Cas9 data, where no effective KO occurs. The wild-type data thus provide a distribution of lineage scores under the background hypothesis (no change upon knockout). For each comparison of populations and each guide library, we first centred and scaled Cas9 data based on the mean and standard deviation calculated from the Non-Cas9 data. The resulting normalised lineage scores are similar to z-scores and were thus used to calculate the probabilities of observing values as extreme (two-sided) using the function *pnorm*. The resulting probabilities can be interpreted similar to p-values, in that they represent the probability of an observed value given a null or background distribution, however with the important difference that in our analyses the null distribution was not obtained based on replicates but based on the distribution of Non-Cas9 data, where no effect is expected. Similar to standard p-values, we next corrected these probabilities for multiple testing using the function *p*.*adjust* with the method “BH” and selected values smaller than 0.05 as significant.

### Validation of single candidates with Flow cytometry

Mouse progenitor cells (LSK) from Cas9 mice were isolated as described above and transduced with CRISPR-seq vectors with NTC or the relevant sgRNAs targeting specific CFs as described above. The cells were cultured *ex vivo* to assess stem cell renewal, lineage priming, myeloid differentiation and terminal myeloid differentiation using the relevant media in every case (see Methods Ex vivo CRISPR screens), harvested and stained with the specific antibody cocktails (see Methods FACS readouts) and analysed with a FACS Aria. FACS data was analysed using FlowJo v10.8.0 Software (BD Biosciences).

### Perturb-seq data generation and sequencing analysis pipeline

#### Library cloning

For each target, we cloned the top 2 sgRNAs identified in the screens in the Lenti-PerturbSeq-BFP vector, which we build from the Lenti-CRISPR-BFP vector (see Extended data Fig 1) by replacing the original sgRNA scaffold for a scaffold that contains the 10X capture sequenced CR1Cs1^53^.

#### *In vivo* Perturb-seq

Primary LSKs were isolated from 8-14 weeks-old ROSA26xCas9 mice following a similar protocol as in the *ex vivo* screens. Then Cas9+ multipotent progenitors were infected with CRISPR libraries to reach 10% infection, which corresponds to a 500X coverage (ratio 50,000 total LSKs per each 10 sgRNAs, where 5,000 cells contain sgRNAs). After transduction, CRISPR modified progenitor cells were grown *ex vivo* for 36 hours under Stem Cell conditions (DMEM/F12, ITSX, PVA, 100 ng/uL Tpo, 10 ng/uL SCF)^54^. Then, cell number and viability were assessed with Cellometer K2 Image Cytometer (Nexcelom Bioscience) and 50,000 viable cells were transplanted to irradiated (902 cGy, 1 minute) 12 weeks-old adult B6.SJL-Ptprca Pepcb/BoyJ (CD45.1) (Jackson #002014) mice via tail injection. Multipotent progenitor cells expand slightly after 36 hours, so, on average we transplanted 1.5 mice for every 10 sgRNAs (delivered to the initial 50.000 LSKs). After two weeks, the transplanted mice were euthanised and bone marrow cells were purified as described for the *ex vivo* screens and stained with BV510 anti Lineage (CD3, CD19, Ter119, CD11b, Gr1) and APC anti cKit antibodies. For single-cell RNAseq we FACS-sorted Lineage- and Lineage+/cKit+ fractions and processed each of them in a 10X single-cell RNA-seq partition aimed at a final coverage of 500 single-cells per CF-KO.

#### *Ex vivo* Perturb-seq

Primary LSKs were isolated from 8-14 weeks-old ROSA26xCas9 mice following a similar protocol as in the *ex vivo* screens. Then Cas9+ multipotent progenitors were infected with CRISPR libraries to reach 30% infection, which corresponds to ∼1,300X coverage (ratio 100,000 total LSKs per each 22 sgRNAs, where 30,000 cells contain sgRNAs). After transduction, CRISPR modified progenitor cells were grown *ex vivo* for 7 days under Stem Cell conditions (see above)^54^ or 2 days under Myeloid differentiation conditions (see above). Then, Cas9+/LV+ cells were FACS sorted for single-cell RNA-seq. 16,000 live cells (cell number and viability assessed with Cellometer K2 Image Cytometer) were processed in a 10X single-cell RNA-seq partition aimed at a final coverage of 1,000 single-cells per CF-KO.

### Perturb-seq in leukaemic cells

0.25-0.5×10^6^ DM Cas9 cells were lentivirally transduced using Retronectin-mediated (Takara Bio #T100A) transduction with pooled guide RNA libraries (Supplemental table 1), and maintained in culture for 6 days post-transduction under standard culture conditions. Transduced, BFP-positive and 7AAD-negative live cells were FACS-sorted (BD Influx; BD Bioscience).

Single-cell RNA-seq libraries were prepared in the Cancer Research UK Cambridge Institute Genomics Core Facility using the following: Chromium Single Cell 3′ Library & Gel Bead Kit v3.1, Chromium Chip G Kit and Chromium Single Cell 3’ Reagent Kits v3.1 User Guide (Manual Part CG000317 for CITEseq and CG000316 for CRISPR). Cell suspensions were loaded on the Chromium instrument with the expectation of collecting gel-beads emulsions containing single cells. RNA from the barcoded cells for each sample was subsequently reverse-transcribed in a C1000 Touch Thermal cycler (Bio-Rad) and all subsequent steps to generate single-cell libraries were performed according to the manufacturer’s protocol with no modifications. cDNA quality and quantity were measured with Agilent TapeStation 4200 (High Sensitivity 5000 ScreenTape) after which 25% of material was used for gene expression library preparation. Library quality was confirmed with Agilent TapeStation 4200 (High Sensitivity D1000 ScreenTape to evaluate library sizes) and Qubit 4.0 Flourometer (ThermoFisher Qubit™ dsDNA HS Assay Kit to evaluate dsDNA quantity). Each sample was normalised and pooled in equal molar concentration. To confirm concentration pool was qPCRed using KAPA Library Quantification Kit on QuantStudio 6 Flex before sequencing. Pools were sequenced on Illumina NovaSeq6000 sequencer with following parameters: 28 bp, read 1; 10 bp, i7 index; 10bp, i5 index and 90 bp, read 2.

#### Chromium 10x single-cell RNA-seq sample data processing

Single cell libraries were generated using the Chromium Next GEM Single Cell 3’ Reagent Kits v3.1 (Dual Index) using the 10x Genomics manufacturer recommended protocol and targeting 16,000 cells per library. The resulting libraries were quantified using qPCR (Roche #4007960166001) and analysed on a Tape Station. Libraries were sequenced in a NextSeq 1000 at 50,000 reads/cell for 3’ Gene Expression libraries and 5,000 reads/cell for CRISPR Feature Barcode libraries.

#### CITE-seq in leukaemia

CITE-seq was performed on 2×10^6^ DM Cas9 murine leukemic cells, stained with Total-Seq B antibodies (Biolegend) for CD11b, Ly6C, CD115, CD14, CD150, CD48, CD34, CD117, CD55, CD41, CD326 and FcgRI (clone M1/70, cat#101273, HK1.4, cat#128053, AFS98, cat#135543, Sa14-2, cat#123341, TC15-12F12.2, cat#115951, HM48-1, cat#103457, SA376A4, cat#152213, 2B8, cat#105849, RIKO-3, cat#131817, MWReg30, cat#133941, G8.8, cat#118247, MAR-1, cat#134341) as per manufacturers protocol. Stained cells were FACS-sorted for 7AAD (BD Bioscience) negative, live cells (BD Influx; BD Biosciences).

### *In vivo, ex vivo* and leukaemia Perturb-seq, trajectory analysis and CITE-Seq data processing and analysis

All analyses in this section were performed in R (version 4.0.2) unless otherwise stated.

#### Basic processing and alignment

Raw reads were processed and aligned to the GRCm38/mm10 reference genome assembly (GENCODE vM23/Ensembl 98) using the software cellranger count (version 6.1.1).

#### Quality control and integration

Initial cell detection from cellranger was used by importing the “filtered” data matrix. Additional quality control and processing was then performed separately for each sample. First, cells with low quality were filtered based on the number of detected genes, UMIs, and the percent of mitochondrial reads using Seurat (version 4.0.0)^55^. For each sample, the 90-th percentile of cells was calculated based on the number of detected UMIs and genes. Cells with less than 20% of either 90-th percentile (and at least 500 genes and 1000 UMIs) were removed. Cells with more than 10% of mitochondrial reads were also removed. Second, cell cycle phases were inferred using the function *CellCycleScoring* from Seurat with default parameters. Third, guide RNAs were assigned to cells using the guide RNA matrix provided by cellranger. In the case of multiple detected guide RNAs in a cell, guides that make up more than 75% of guide reads in this cell were used. Fourth, data were aligned across samples using the function *align_cds* from Monocle3 (version 0.2.3.0)^56^, selecting the sample as the alignment group and the cell cycle phase as the residual model. Finally, the UMAP projection and clustering was calculated using the functions *reduce_dimension* and *cluster_cells* from Monocle3 on the aligned data.

#### Cell type assignment (in vivo and ex vivo)

Cell types were predicted using the package singleR (version 1.4.1), based on a dataset obtained from Izzo and colleagues^48^ and a dataset of bone marrow obtained from the packages CytoTRACE (version 0.3.3)^57^. SingleR^58^ was run using the Wilcoxon method for differential analysis. Cells in clusters with more than 80% of cells predicted as granulocytes, granulocytes progenitors, or immature B cells in the bone marrow dataset from CytoTRACE were assigned based on this dataset. All other clusters were assigned based on the predictions by Izzo et al^48^. Eosinophils and basophils were combined in one label. Cells predicted as erythrocytes were further split into MEPs, Erythroid Progenitors or Erythrocytes based on a comparison of gene signatures with external datasets^49^ and on the expression of key marker genes: 1) Low *Gata2*/High *Gata1, Epor* & *Klf1* marked the transition from MEPs to Erythroid progenitors 2) Induction of *Hba-a1* and *Hbb-b1*, increased *Tfrc* expression, enrichment of S and G2 cell-cycle signatures mark the transition from Erythroid Progenitors to Erythrocytes.

Next, cell constituting less than 10% of a given cluster were re-assigned to the cell type representing the majority in each cluster. Then, MEPs with *Gata2* expression greater than *Gata1* expression were labelled as early MEPs. Also, MEPs clusters with strong cell cycle phase signatures were labelled accordingly. A cluster of MEPs with a large fraction of cells harbouring *Rcor1* guides was labelled as “MEP (perturbed)”.

#### Cell type enrichment analyses

To test differences in distributions of knockouts (KOs) and non-targeting controls (NTCs) across the identified cell types, we tested the enrichment of Kos compared to NTCs within each cell type compared to all other cell types. We first tested enrichment in broad cell types (HSCs, myeloid cells, megakaryocytes, basophils, erythrocytes, and lymphoid cells). Second, we tested enrichment only between myeloid and erythroid cells. Clusters with less than 5 NTCs or less than 25% of NTCs were removed from this analysis. The Fisher Exact test was used with the function *fisher*.*test*. Enrichment was tested against each NTC separately.

*Cross-projection of ex vivo and leukaemia samples to in vivo data*. To project *ex vivo* and leukaemia samples onto the *in vivo* data, we adapted the ProjecTILs algorithm (version 2.0.2)^59^, predicting UMAP coordinates for each cell from the ex vivo and leukaemia samples based on the UMAP coordinates of *in vivo* cells using a k-nearest-neighbour approach with k set to 20 neighbours.

#### Differential expression analysis

To identify molecular effects of CF-KOs, we performed differential expression comparing cells with CF-KOs to NTCs using the nebula package (version 1.1.8), which implements a fast negative binomial mixed model. We removed cluster with less than 31 cells and genes with less than 21 reads. We then ran nebula^60^ with default parameters, testing differences of KOs to NTCs with fixed effects (parameter “pred”) and adding sample information as random effects (parameter “id”). P-values were corrected for multiple testing using the function *p*.*adjust* with the method “BH”. Next, gene set enrichment was performed using the function *fgsea* from the *fgsea* package^61^.

#### Pseudotime trajectory analysis

Pseudotime for the different lineage trajectories were identified using diffusion maps^62^ applied to the non-targeting control population, using Scanpy (version 1.9.1)^63^ functions *tl*.*diffmap* and *tl*.*dpt*. Perturbed cells were then mapped to their nearest k=15 non-targeting cells in PCA space, considering the first N=8 principal components, and then assigned the mean pseudotime value across these cells. Principal component analysis cut offs were found via elbow plots through assessing the variance accounted for in the first N principal components. Each branch was extracted for separate analysis using the aforementioned cell labels (see Figure 2a) and pseudotime was scaled to the unit interval. We plot perturbations with crucial biological significance, which incidentally show visually striking distribution differences compared to the non-targeting cell population.

Cite-seq read counts obtained from cellranger were normalized to log counts per million (CPMs) and then scaled.Perturb-seq data in leukaemia were processed in the same way as the in vivo data, as described above. Enrichment analyses were performed on clusters instead of cell types.

### CF-KO Chromatin accessibility profiling

#### Isolation of progenitors, CRISPR loss of function and *ex vivo* differentiation

For each reaction we FACS-sorted 20,000 haematopoietic progenitor cells (LSKs) from the Cas9 strain as outlined above. Then we transduced them with Lenti-CRISPR-BFP virus, containing the top performing sgRNA against each CF and cultured the cells for 48h under multipotent conditions detailed above (Tpo, SCF, PVA). After that, the CRISPR-modified progenitors were stimulated with cytokine cocktails for lineage-priming or myeloid differentiation for 4 days (see media composition above). Finally, the CRISPR edited progeny was FACS-Sorted (BFP+) and harvested by centrifugation for ATAC-seq.

#### Fast ATAC-seq

Freshly sorted cells were collected in 1X PBS + 0.05% BSA, spun at 300G for 12 min at 4 ºC and resuspended in 20 uL transposition mix: 1X TD buffer and 2 uL of TDE1 (Illumina #FC-121-1030), 0.01% Digitonin in DMSO, 0.1% Tween20, 0.1% IGEPAL in Molecular Biology Grade water. Tagmentation was performed at 37 ºC for 30 minutes in an Eppendorf ThermoMixer with agitation at 1000 rpm, and stopped by adding EDTA to a final concentration of 5 mM. Then, 2 uL of Proteinase K were added and the samples were incubated for 30 minutes at 40 ºC. Genomic DNA was purified with a 2X SPRI cleanup and tagmented sites were PCR-amplified with 25 uL Kappa HIFI 2X and 2.5 uL of 10 uM P5-i5-read1 and P7-i7-Read2 primers (Suppl Table 3). The PCR cycling conditions are: 5 min at 72 ºC, 10 x [2 min at 98 ºC, 20 s at 98 ºC, 30s at 60 ºC, 1 minute at 72 ºC] 3 min at 72 ºC. Amplified ATAC-seq libraries were purified with a 1.6X SPRI Cleanup, quantified using the Qubit dsDNA high sensitivity assay kit, and then analysed on a Tape Station 4000. ATAC-seq libraries were sequenced at 50 million reads on a NextSeq 1000.

#### ATAC-Seq data processing and analysis

Based on the ATAC-seq nf-core pipeline^64^, we ran Trim Galore (version 0.6.6)^65^ with Cutadapt (version 3.4)^66^ using the default parameters to trim low-quality and adaptor sequences in paired ATAC-seq reads. We then aligned these reads to the GRCm38/mm10 reference genome assembly using Bowtie2 (version 2.3.4.2)^67^ with parameters - X 1000 --no-discordant --no-mixed --very-sensitive. Subsequently, we removed duplicated regions with Picard (version 2.25.4) (Broad Institute, G. R. Picard Toolkit. https://broadinstitute.github.io/picard/ (2019)), non-interesting chromosomes (chrM, chrUn, …), and blacklist regions included in the ENCODE blacklist (version 2.0)^68^. Finally, we removed Tn5 adaptors with alignmentSieve’s (version 3.5.1)^69^ –ATACshift parameter and indexed the final BAM files with SAMtools (version 1.3.1)^70^. These BAM files were then processed to CPM scaled Bigwig files with bamCoverage (version 3.5.1)^71^.

To identify the ATAC peaks for each sample, we pooled replicates and converted the paired BAM files to single-read bed format using function bamToBed from BEDTools (version 2.27.1)^70^. Then, we used MACS (version 2.2.7.1)^72^ with parameters –broad -f BED --keep-dup all -- nomodel --shift −75 --extsize 150 to call peaks. To compare peak strength between conditions, we generated a unified peak set for all experiments, ending with 376,658 and 207,724 peaks in LSKs and Npm1c/Flt3-ITD murine leukemic cells (DM-Double Mutant), respectively. We then annotated these consensus peaks with the function *annotatePeaks* from HOMER (version 4.10)^73^, counted the reads on them with featureCounts (version 2.0.1)^74^, and calculated adjusted CPM values with edgeR’s (version 3.34.1) TMM method^51^. Additionally, we used DESeq2 (version 1.32.0)^75^ to measure the Fold Change (FC) between conditions, defining the peaks with absolute log2(FC) values greater than 0.75 and adjusted p-values lower than 0.01 as differentially enriched. Finally, we filtered out peaks with less than 2 CPM and/or 10 reads in the compared two conditions (KO vs. WT).

#### Motif analysis in ATAC-seq peaks

To look for differential TF motifs enrichment between KO and WT, we used TOBIAS (version 0.13.2)^26^. Following the program guidelines, we generated a consensus set of peaks with all the peaks called previously in both the control and the compared KO sample. We then renamed and formatted HOMER’s list of vertebrate known.motifs to make it suitable for TOBIAS. After generating the TF footprint bigwigs (using the function *ATACorrect* with parameters --read_shift 0 0 and the function *ScoreBigwig* with default parameters), we computed the differentially bound motifs with the function *BINDetect*.

Additionally, we used the output TF binding coordinates of each of the motifs to measure the gain/loss of TFs union to chromatin for each chromatin factor KO. Then, we compared these gain/loss coordinates between different KOs to measure their degree of overlap.

### ChIP-seq analysis of healthy and leukemic populations

#### Isolation of hematopoietic cells from bone marrow

Murine haematopoietic cells were harvested for the isolation of GMP and MEP progenitor cells and myeloid, erythroid and B-cell lineages from bone marrows of 12 weeks-old C57BL6 mice.

Each cell type was FACS-sorted into PBS according to the following strategy.

GMP: Lineage (CD3, CD19, CD11b, Gr1, Ter119, B220)-, cKit+, Sca-1-, FcgRIII+, CD34+

MEP: Lineage (CD3, CD19, CD11b, Gr1, Ter119, B220)-, cKit+, Sca-1-, FcgRIII-, CD34-

Monocytes: CD3-, CD19-, Ter119-, CD11b+

B-cell: CD3-, CD19+, Ter119-, CD11b-

Erythroid cells were FACS-sorted from spleens of 12 weeks-old C57BL6mice as: CD3-, CD19-, Ter119+, CD11b-, Gr1-.

After sorting, cells were pelleted at 500xG for 7 min with low brake.

#### Isolation of leukaemic cells

*Npm1c/Flt3-ITD/Cas9* double mutant (DM) cells were generated from lineage-depleted, bone marrow cells of primary transgenic mice post-leukemic onset as previously described^76,77^. Cells were maintained in XVIVO-20 medium () supplemented with 5% Fetal Bovine Serum (FBS) (ThermoFisherScientific #10270106), 1% PSG (Gibco #10378-016) and recombinant murine SCF 50 ng/mL (PeproTech # 250-03-250UG), murine IL-3 10 ng/mL (PeproTech #313-13-10UG) and murine IL-6 10 ng/mL (R&D Systems #406-ML-025), in a 37°C and 5% CO_2_ atmospheric environment.

#### Crosslinking

Freshly sorted normal cell or early passage leukaemic cells were pelleted as described above, resuspended and rocked in a 5 mL solution of 3 mM EGS, DSG, and DMA crosslinkers in 1X PBS for 20 min at room temperature followed by 5 minutes with 1% formaldehyde. The crosslinking was quenched by adding Glycine to 125 mM final concentration. Then cells were pelleted, washed twice with cold 0.5% BSA in PBS containing 1X Protease Inhibitors (Roche) and flash frozen at −80C.

#### Chromatin Immuno-precipitation

Crosslinked cells were incubated in Cell Lysis Buffer (10 mM HEPES pH 7.5, 10 mM NaCl, 0.2% IGEPAL) + Protease Inhibitors (Roche) for 5 minutes on ice, nuclei were pelleted at 5,000 g, resuspended in Sonication buffer (0.5% SDS, 5mM EDTA) and pelleted again at 8,000 g, which yielded a translucent chromatin pellet. This pellet was resuspended in 50-100 uL of Sonication buffer and sonicated for 5 cycles (30s ON, 30s OFF) in a Bioruptor Nano.

Then 4 volumes of ChIP dilution buffer (25 mM HEPES, 185 mM NaCl, 1.25% Tx-100) + Protease Inhibitors were added to each sample, vortexed and centrifuged at 20,000 g for 10 minutes. The supernatant, containing the solubilised chromatin fragments, was transferred to Protein low bind tube and incubated for 10-12 hours with the relevant antibodies (see below). Then, 25 uL Magna ChIP Prot A+G were added and incubated for 2h at 4C to capture the Immune-complexes, which were then washed 2X with RIPA, 2X with RIPA-500, 2X with LiCl buffer and 1X with TE buffer. ChIPed DNA was released and reversed crosslinking by 45 min incubation with 2 uL of Proteinase K in ChIP Elution buffer followed by 1-hour incubation at 68 ºC. Finally, the ChIPed DNA was purified with a 2.2X SPRI Cleanup and quantified by Qbit.

#### Preparation of ChIP-seq libraries

ChIP-seq libraries were prepared from 1-10 ng of DNA with the NEB Next Ultra II kit with the following modifications:

- End Prep: the reaction was performed in half of the volume specified in the manual with the following program.
- Adaptor Ligation: the reaction was performed in half of the volume specified in the manual with the following program.
- Final PCR cycles were adjusted to the initial input DNA amount according to the NEB Next Ultra II kit protocol.

ChIP-seq libraries were QCed for concentration (Qbit) and size (Tape-Station) and sequenced to 100 million reads per sample.

#### ChIP-seq data processing and analysis

Based on the ChIP-seq nf-core pipeline^63^ we first processed the FASTQ files to BAM files as described in the **ATAC-Seq data processing and analysis** section (skipping the Tn5 adapter removal). Next, to identify peaks for each sample, we pooled replicates and used MACS^71^ with parameters -f BAMPE --keep-dup all. To compare peak strength between cell types, we generated a unified peak set per CF (Brd9, Kmt2a, Kmt2d, and Smarcb1). We then followed the steps explained in the **ATAC-Seq data processing and analysis** section to annotate the peaks and calculate the CPM reads on them. Finally, we measured the FC between cell types in one of two ways, based on the availability of replicates: In sample pairs with replicates, we used DESeq2^75^ to get peaks with an absolute log2(FC) greater than 0.75 and an adjusted p-value lower than 0.01. In sample pairs without replicates, we set a threshold of log2(FC) greater than 1. Additionally, we filtered out peaks with less than 2 CPM and/or 10 reads in the compared two conditions and with a FC of CF/IgG lower than -1.

#### Motif analysis in ChIP-seq peaks

We first generated a list of cell-type-specific peak coordinates for each of the analysed CFs (Brd9, Kmt2a, Kmt2d, and Smarcb1). To do so, we selected all peaks that were significantly enriched or depleted in pairwise comparisons of GMP, Myeloid, MEP, Ery, and B-cell on each of the CFs. Then we clustered and manually curated these coordinates to get a list of cell-type-specific peaks per CF. Enrichment of transcription factors in cell-type-specific peak coordinates was then analysed using the function *findMotifsGenome* from HOMER^72^. For each CF, cell-type specific peaks were compared to all peaks found across all five cell types and all four CFs as background. Motif enrichment analyses were centred on the 100 bp surrounding the peak summit.

#### Comparison of normal and leukaemic patterns

To identify transcription factor switches in leukaemia, we defined subsets of peaks that were gained in DM AML (Leukaemic), shared in DM and GMP (Common), and not present in DM but present in GMP and Monocytes (Normal). Gained peaks were defined by log2(FC) values greater than 1, lost peaks by log2(FC) values smaller than −1, and shared peaks by absolute log2(FC) values lower than 0.5. The subsetting was done per CF in a consensus peak dataset of Smarcb1, Kmt2a, and Kmt2d. Similar to **Motif analysis in ChIP-seq peaks** section, we looked for enrichment of transcription factors in each of the subsets using the function *findMotifsGenome* from HOMER, and all peaks from the consensus (across all subsets) were used as the background set.

To measure Stat5a binding signal over the CF-bound sites we collected Stat5a ChIP-seq sequencing data at the same consensus coordinates stated right above and computed the CPM values as described in **ATAC-Seq data processing and analysis** section.

### AML single guide functional assays

*Cell transduction and sorting of differentiated populations*. Npm1/Flt3-ITD/Cas 9 DM murine AML cells were lentivirally-transduced using a Retronectin-transduction protocol (T100A; Takara Bio) with pHKO9 containing guide RNAs complementary to murine BAF complex members (*Smarcb1, Smarcd2, Brd9* and *Smarcd1*), COMPASS complex members (*Kmt2a, Kmt2d* and *Wdr82*), Repressors (*Hdac3* and *Setdb1*), *Stat5a*, or a corresponding non-targeting control (NTC). A single guide RNA was selected for each gene target.

Transduction was confirmed at 2 days post-transduction by measurement of percent BFP-positive (BFP+) cells by flow cytometry. Dropout was monitored by flow cytometric assessment (BD LSR Fortessa II; BD Bioscience) of BFP+ cells in culture at 6 days, 9 days and 13 days post-transduction. Fold change in BFP+ cells at each time point relative to the proportion of BFP+ cells at day 2 is presented. Immunophenotypic analysis was also performed by flow cytometry (BD LSR Fortessa II; BD Bioscience) at 6 days post-transduction with guide RNAs for *Smarcb1, Smarcd2, Kmt2a* and NTC. Briefly, transduced DM cells were stained for CD11b (PE-Cy7 conjugated; clone M1/70; BD Biosciences), Gr-1 (Ly6G/Ly6C; APC-Cy7 conjugated; RB6-8C5 clone; BD Biosciences) and CD55 (PE-conjugated; clone RIKO-3; Biolegend). Percent Gran-like (CD11b-high/Gr-1+) and Ery/Baso-like (CD55-high) specifically in the transduced, BFP+ subset was assessed. All flow cytometry data were analysed with FlowJo (Tree Star; v10.8.1). P-values were calculated using a two-way ANOVA statistical test or ratio-paired t-test (Prism; GraphPad software; v9.1).

#### Clonogenic and Cell proliferation assays of DM AML cells

2×10^6^ DM murine leukemic cells were stained for CD11b (PE-Cy7 conjugated; clone M1/70; BD Biosciences), Gr-1 (Ly6G/Ly6C; APC-Cy7 conjugated; RB6-8C5 clone; BD Biosciences), CD55 (PE-conjugated; clone RIKO-3; Biolegend), CD41 (APC-conjugated; clone MWReg30; Biolegend) and CD34 (FITC-conjugated; clone RAM34; BD Bioscience). Gran-like (CD11b-high/Gr-1+); Ery/Baso-like (CD55-high/CD41-) and CD34+ fractions were subsequently FACS-sorted (BD Influx; BD Bioscience). 1000 cells of each sorted population were seeded in 1 mL of methylcellulose medium supplemented with recombinant murine SCF and IL-3 with recombinant human IL-6 and Epo (M3434, STEMCELL Technologies) in duplicate. Methylcellulose cultures were maintained in a 37 °C and 5 % CO_2_ atmosphered environment, and total colony forming units (CFU) enumerated 7 days later. Photographs of colonies were obtained on the STEMvision instrument and software (STEMCELL Technologies). Data is presented as a fold change of average CFU per 1000 cells seeded, relative to the CD34+ fraction. 10^4^ Gran-like, Ery/Baso-like or CD34+ DM cells were also maintained in standard culture conditions for 21 days, and number of cells counted every 7 days. Total cell number is presented as a fold to corresponding CD34+ counterpart for each timepoint. P-values were calculated using a two-way ANOVA statistical test (Prism; GraphPad software; v9.1).

#### Ethical compliance

Murine ethical compliance was fulfilled under the Guidelines of the Care and Use of Laboratory Animals and were approved by the Institutional Animal Care and Use Committees at University of Navarra, Spain, and the Animal Welfare Ethical Review Body at the University of Cambridge, UK. Research in the UK was conducted under Home Office licence PP3042348.

#### Data and code availability

The data used in and generated from this publication will be made available at the time of publication. Code is available at https://github.com/csbg/tfcf.

## Additional data

**Supplementary Table 4** - Products (Media, cytokines, etc) with catalogue number refs

**Extended Data Figure 1** – Map of the Perturb-Seq vector

## Acknowledgements

Research in B.J.P.H’s laboratory was supported by Cancer Research UK (C18680/A25508), the European Research Council (647685) and the Cancer Research UK Cambridge Major Centre (C49940/A25117). D.L.-A. is a Marie-Skłodowska-Curie International Fellow (886474) and is supported by a Junior Research Grant from the European Hematology Association. N.N. is supported by a Kay Kendall Leukaemia Fund Junior Fellowship (KKL1440). C.L. is supported by a Junior Research Grant from the European Hematology Association. The project leading to these results has received funding from “la Caixa” Foundation under the agreement LCF/PR/HR20/52400016. This research was supported by the NIHR Cambridge Biomedical Research Centre (BRC-1215-20014). The views expressed are those of the authors and not necessarily those of the NIHR or the Department of Health and Social Care. This research was funded in part, by the Wellcome Trust (203151/Z/16/Z) and the UKRI Medical Research Council (MC_PC_17230) who supported the Wellcome – MRC Cambridge Stem Cell Institute. For the purpose of Open Access, the author has applied a CC BY public copyright licence to any Author Accepted Manuscript version arising from this submission.

## Author contributions

D.L.-A., N.F. and B.J.P.H. designed the study. D.L.-A. and B.J.P.H. wrote the manuscript. D.L.-A., A.G.-S., J.M., T.G., J.T.-K., F.P., N.F. and B.J.P.H. analysed data. D.L.-A., A.G.-S., N.N., C. D. V., G.G., M.N., J.Z., F.M., N.T., I.C., C.L., D.A., A.L. and B.S.-O. performed experiments and collected data. B.J.P.H. supervised the study.

