## Supplementary figures and images for "Systematic functional screening of chromatin factors identifies strong lineage and disease dependencies in normal and malignant haematopoiesis"

### Extended Data Figure 1

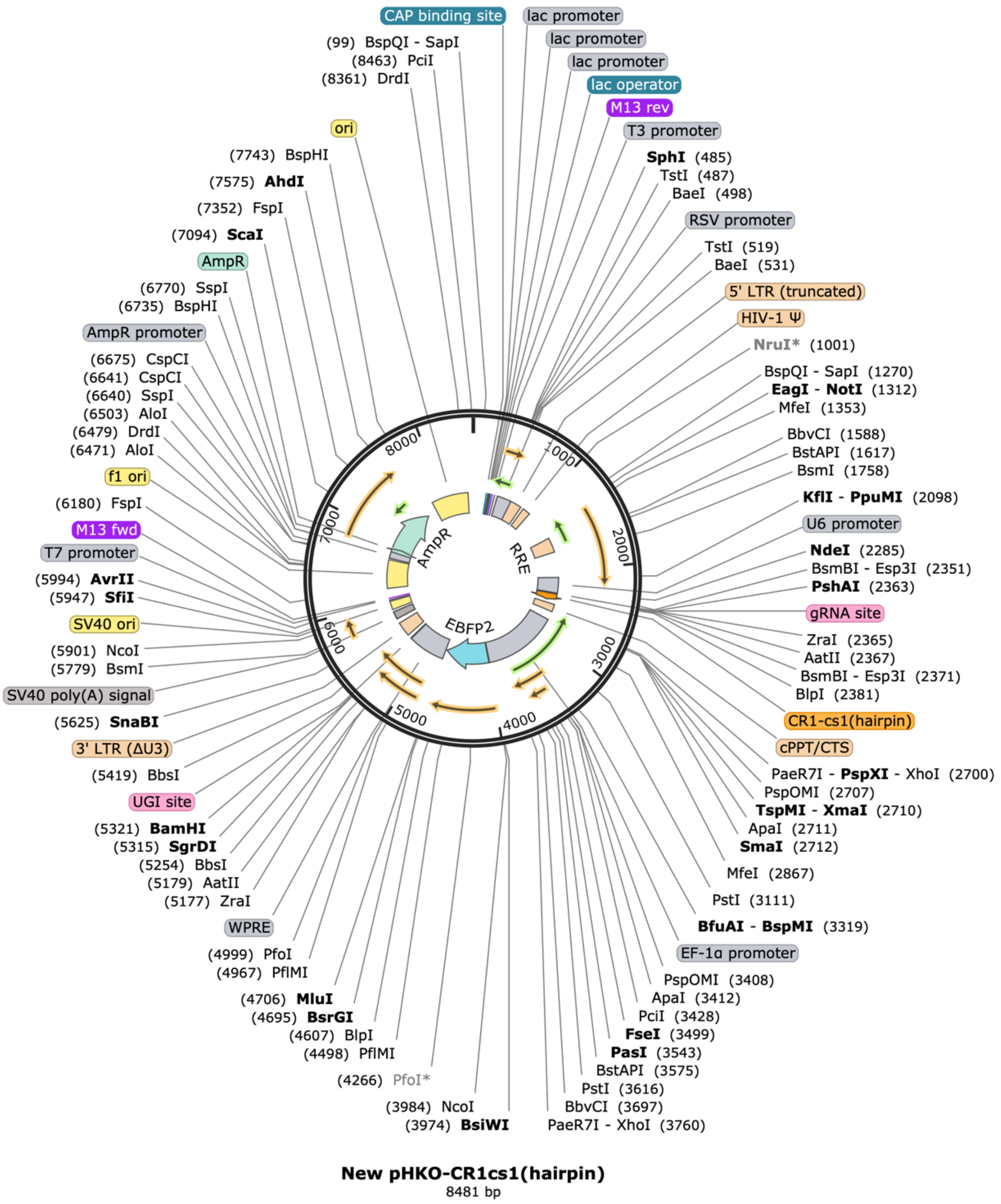


Lenti-CRISPR-BFP vector

### Supplemental Figures

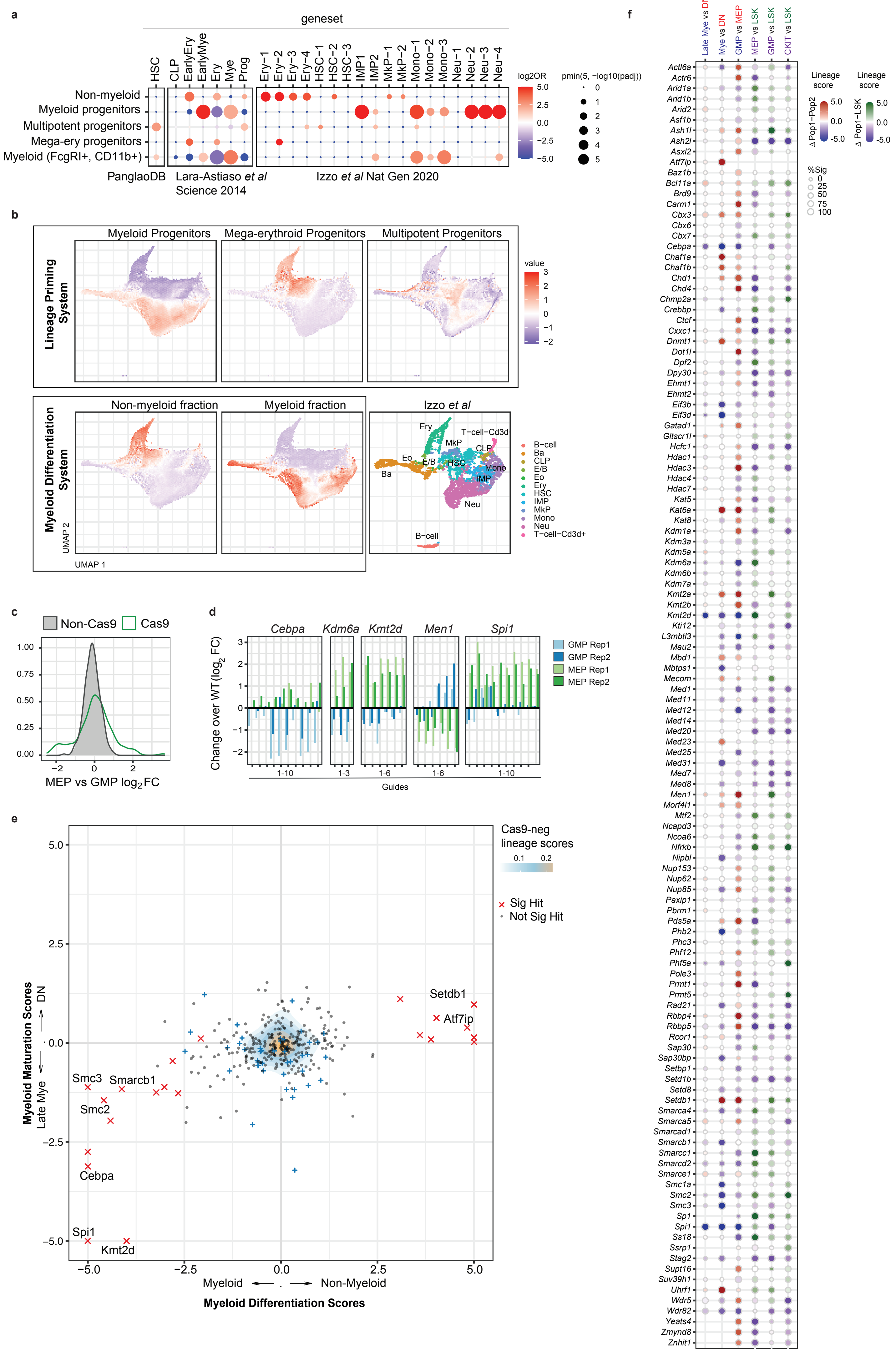

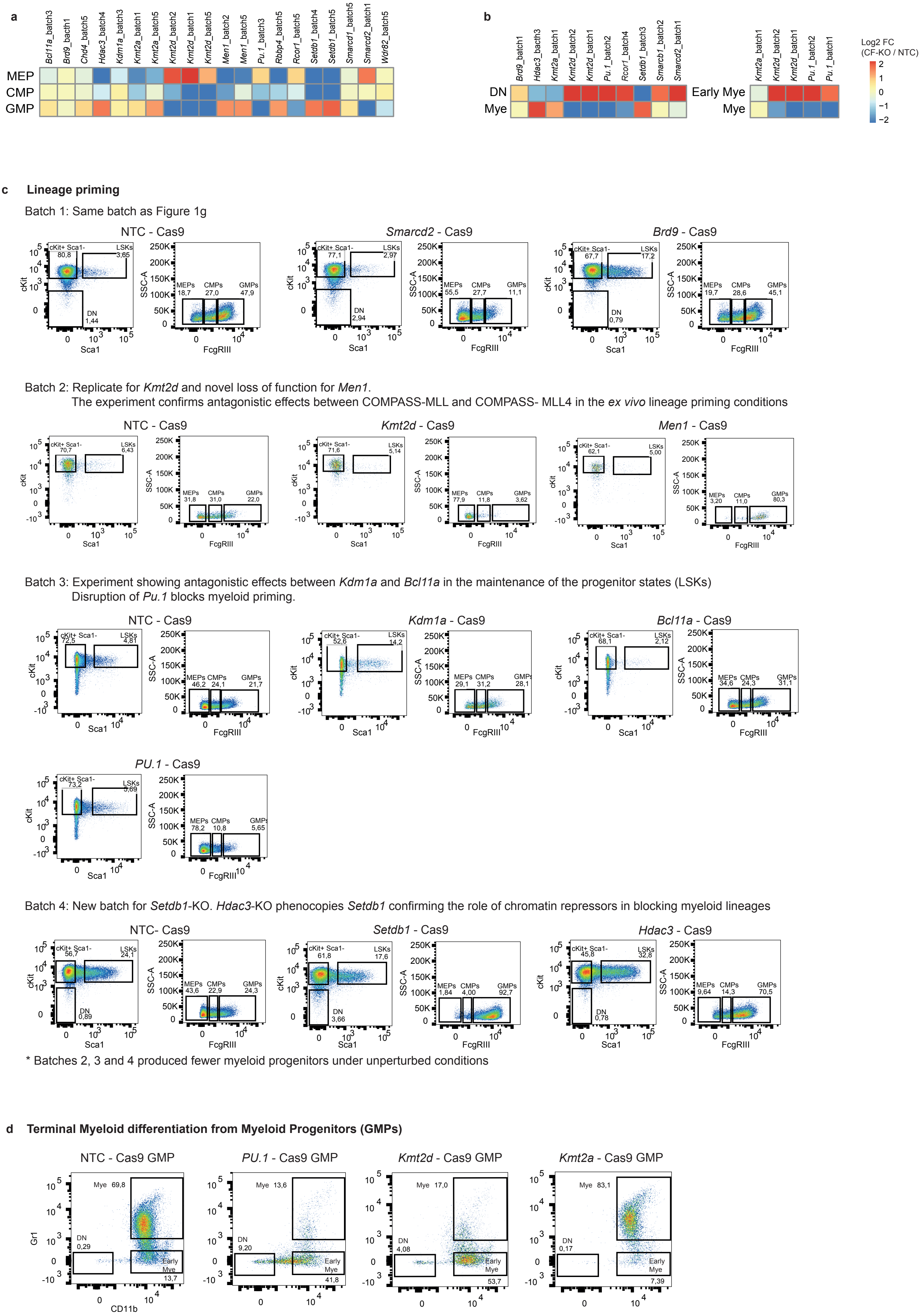

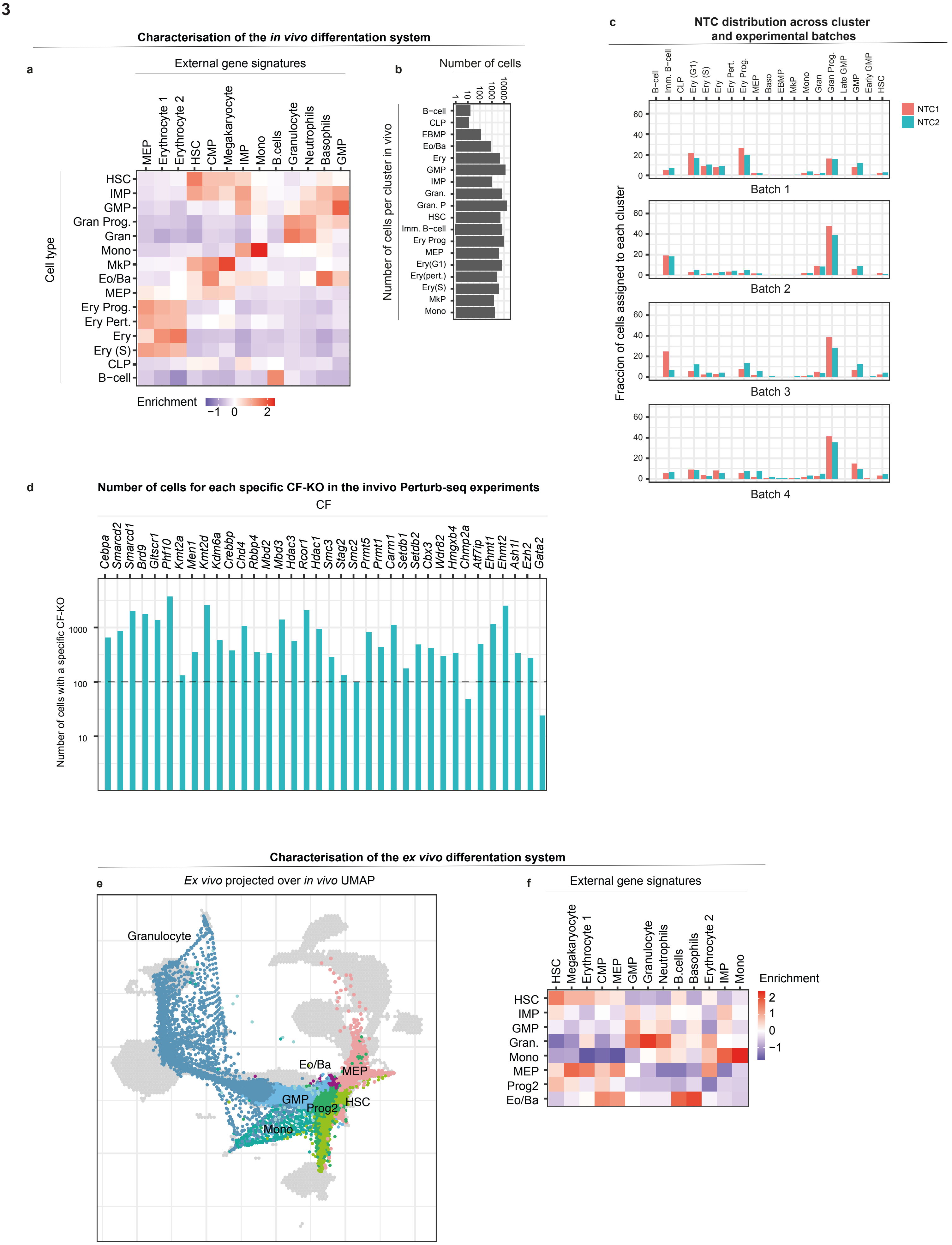

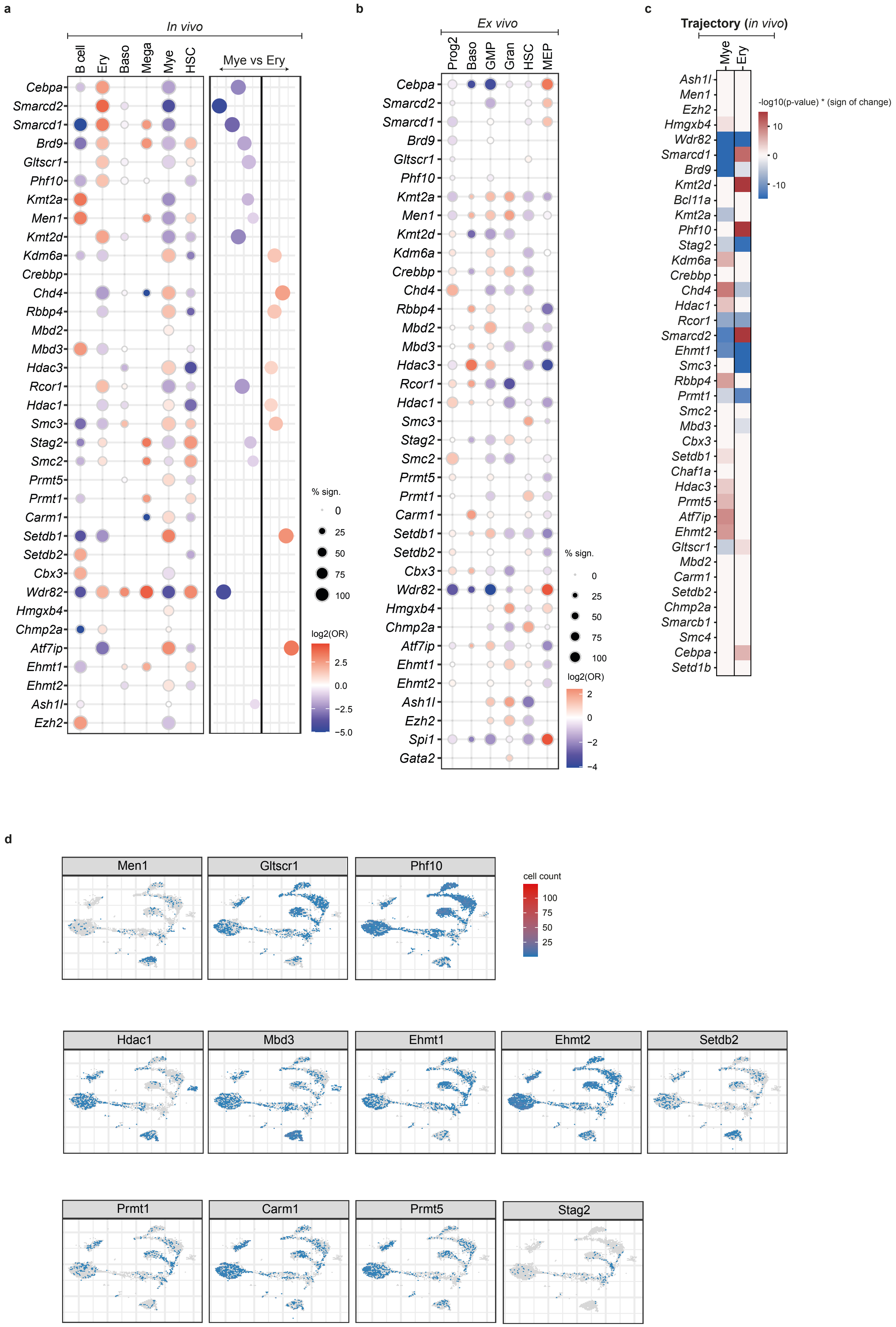

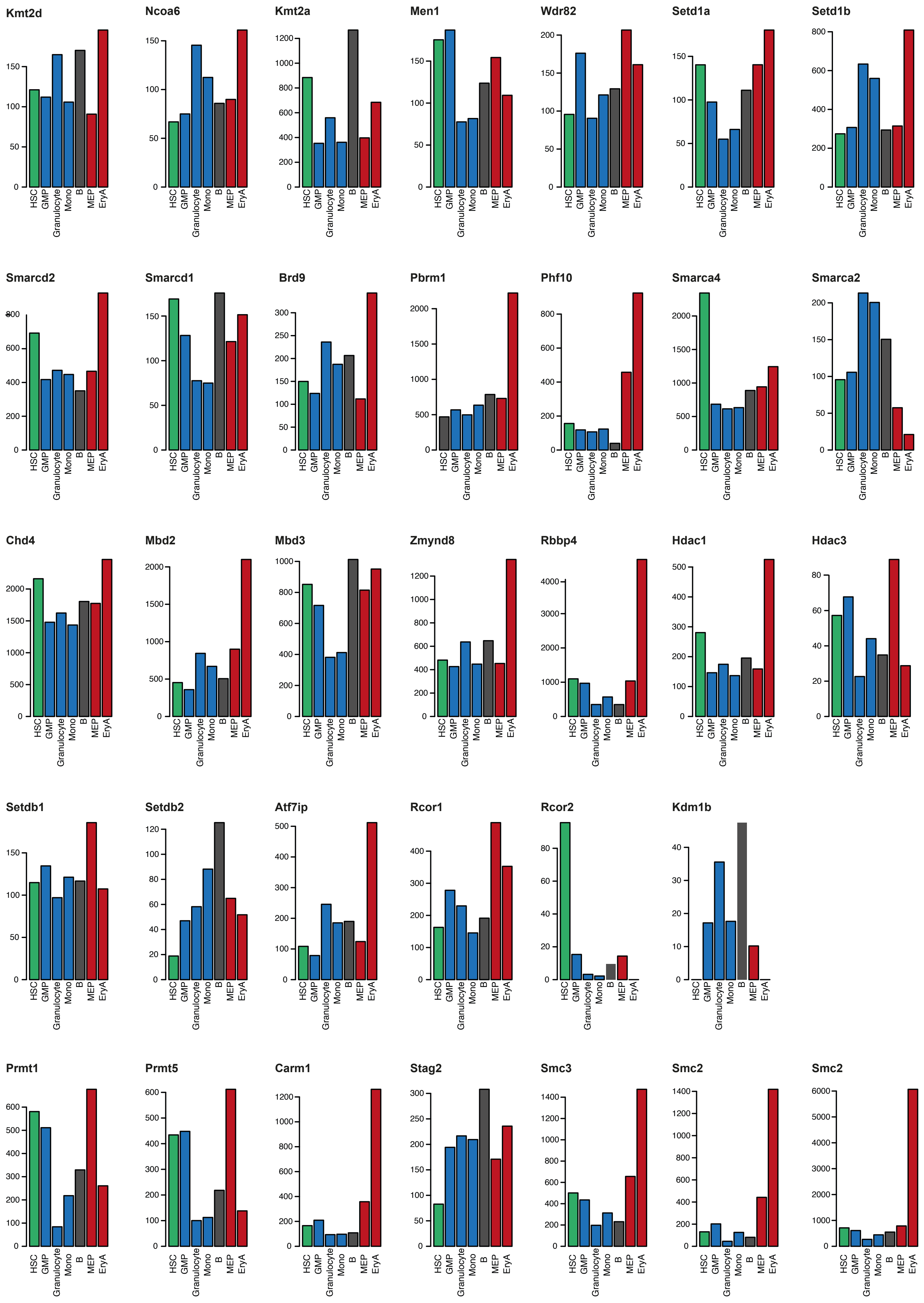

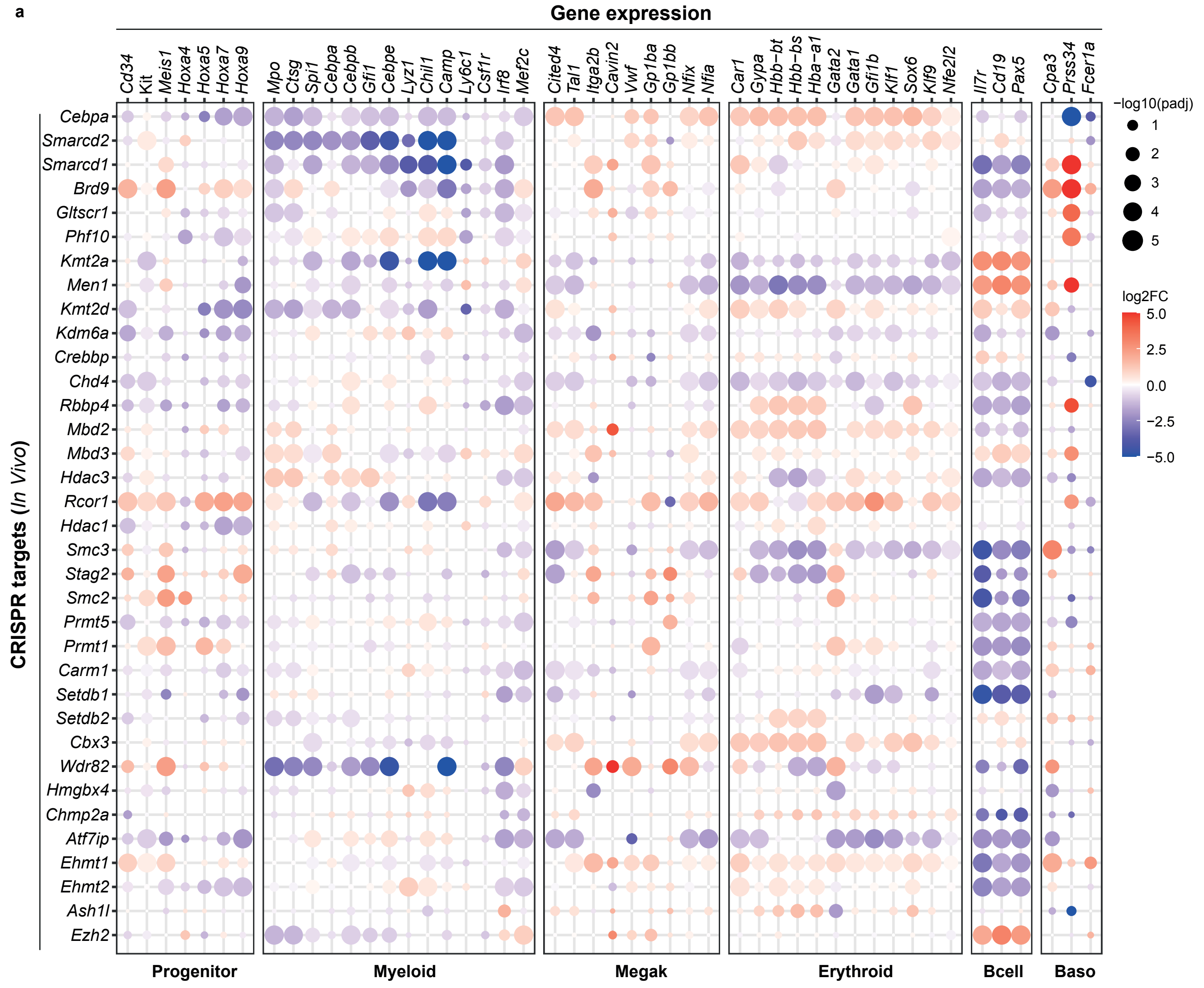

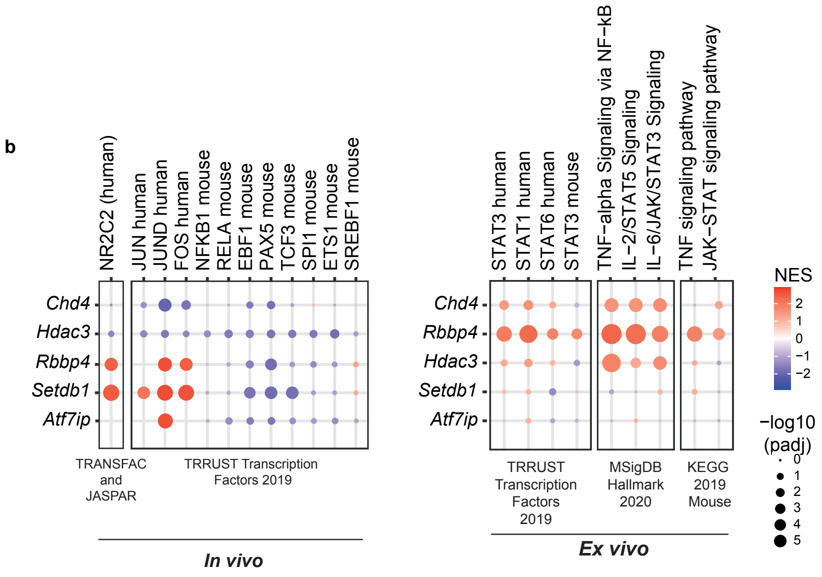

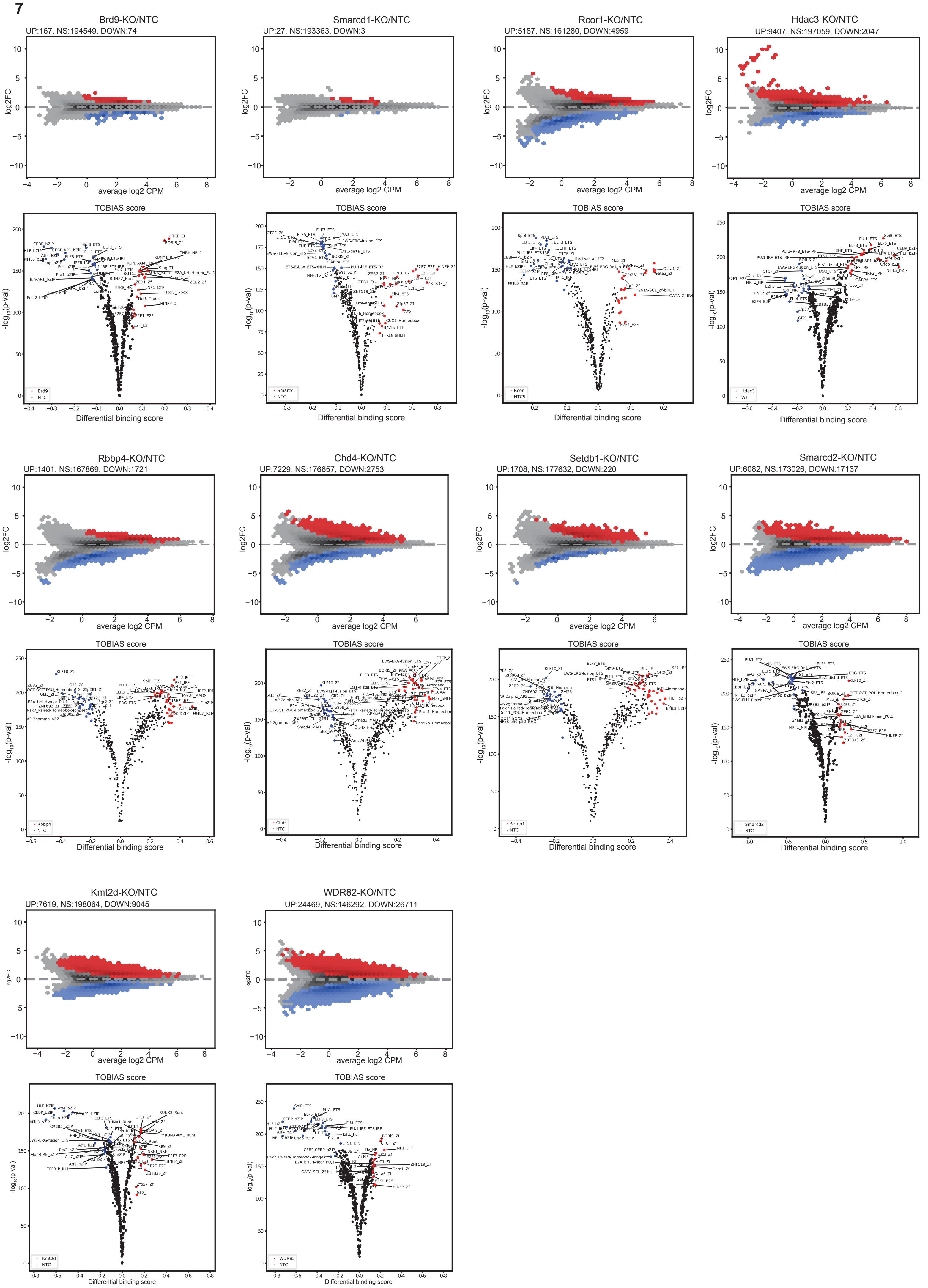

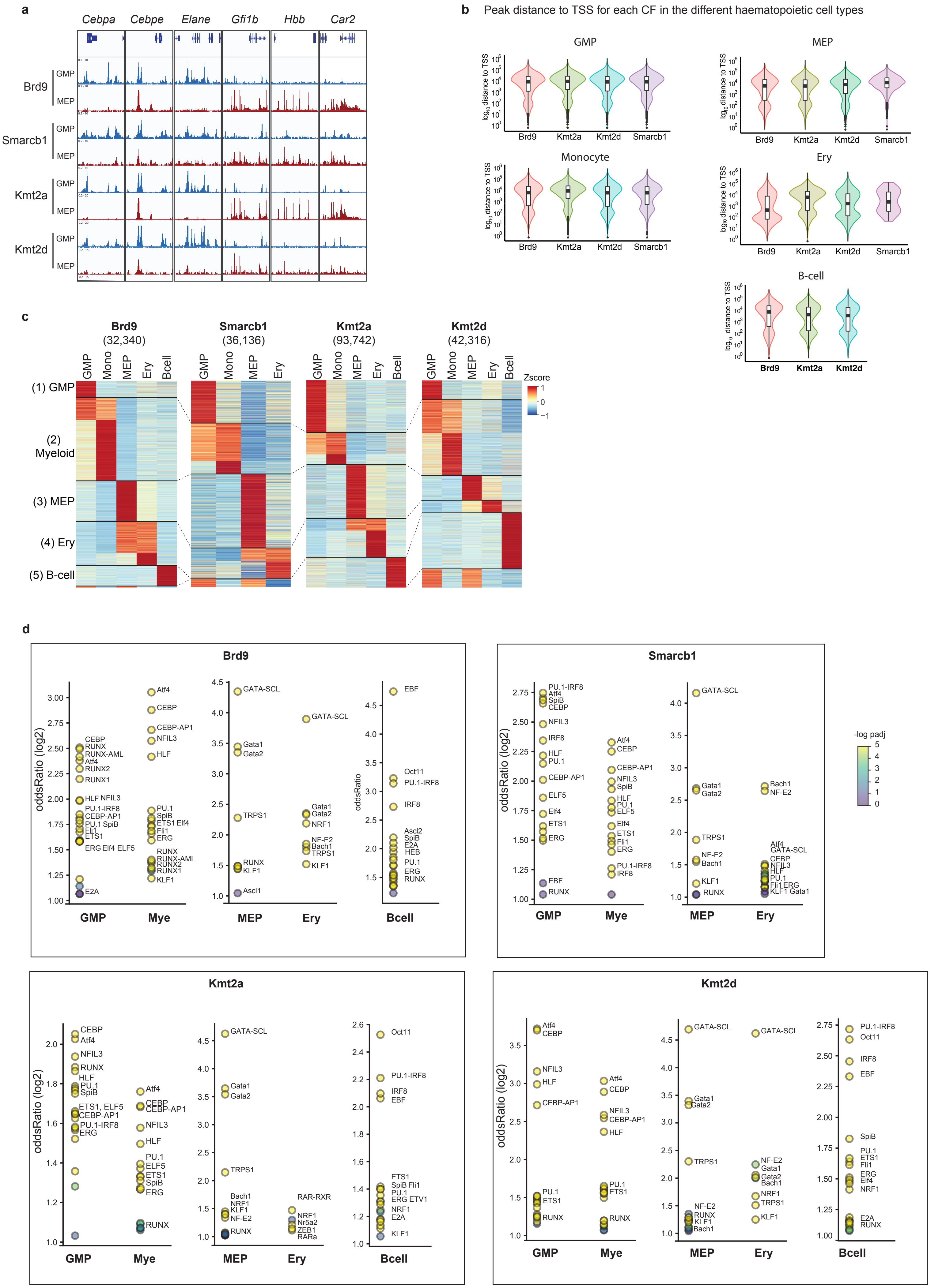

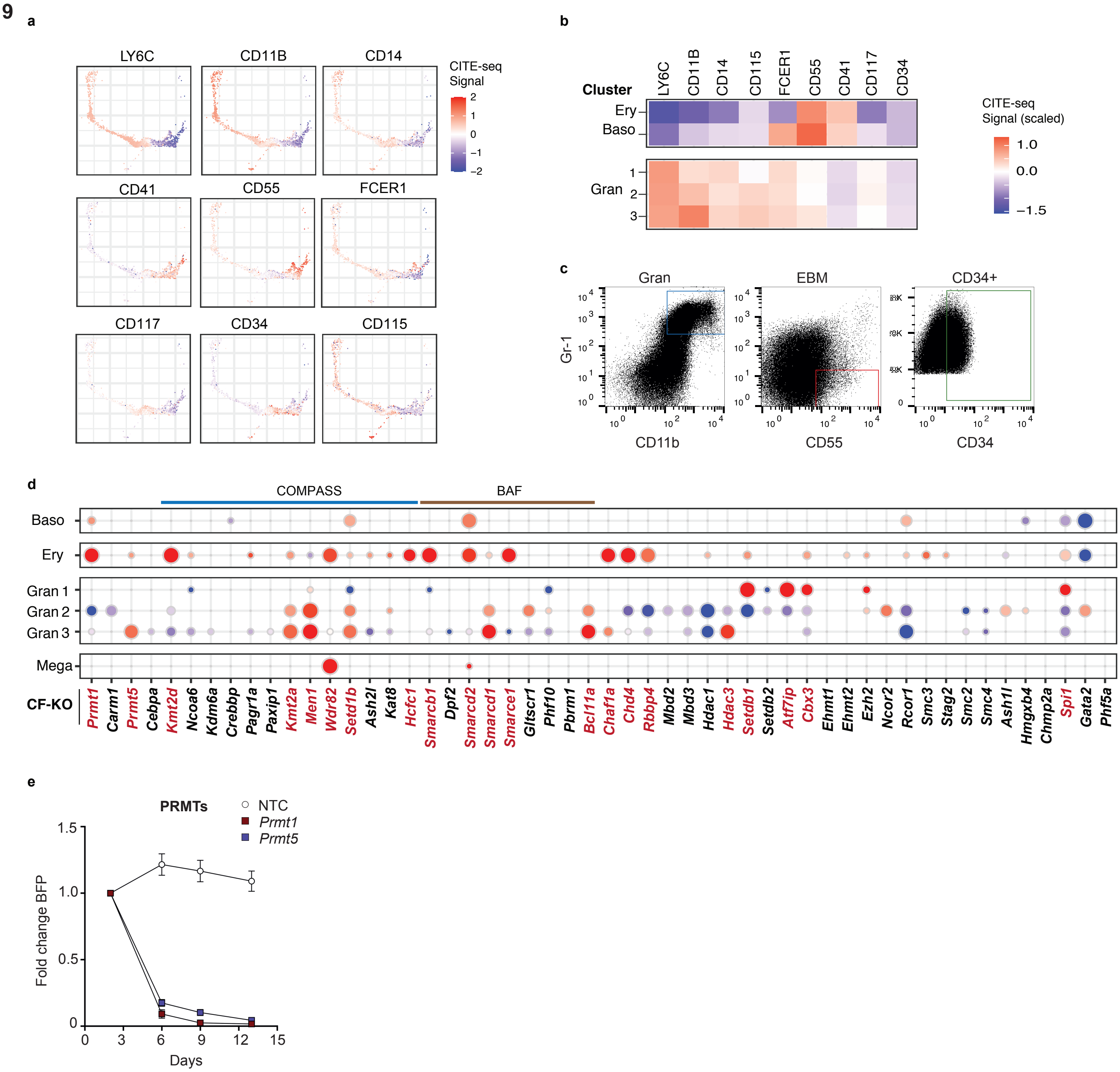

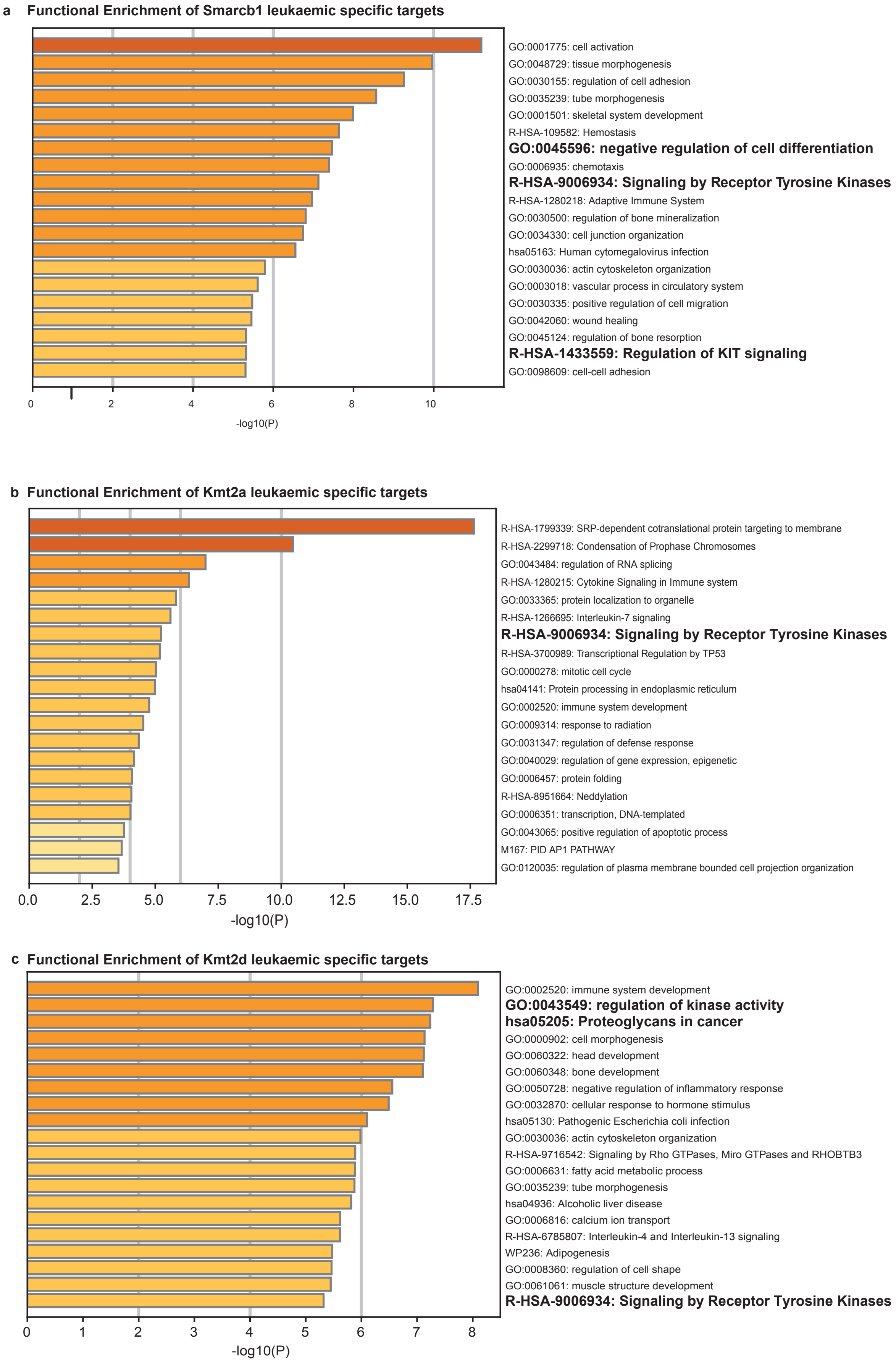
